# A general framework for straightforward model construction of multi-component thermodynamic equilibrium systems

**DOI:** 10.1101/2021.11.18.469126

**Authors:** Nick H. J. Geertjens, Pim J. de Vink, Tim Wezeman, Albert J. Markvoort, Luc Brunsveld

## Abstract

Mathematical modelling of molecular systems helps elucidating complex phenomena in (bio)chemistry. However, equilibrium conditions in systems consisting of more than two components can typically not be analytically determined without assumptions and resulting (semi-)numerical models are not trivial to derive by the non-expert. Here we present a *framework* for equilibrium models that utilizes a general derivation method capable of generating custom models for complex molecular systems, based on the simple, reversible reactions describing these systems. Several molecular systems are revisited via the *framework* and demonstrate the simplicity, the generality and validity of the approach. The ease of use of the *framework* and the ability to both analyze systems and gain additional insights in the underlying parameters strongly aids the analysis and understanding of molecular equilibrium systems. This conceptual *framework* severely reduces the time and expertise requirements which currently impede the broad integration of these highly valuable models into chemical research.

---

Complexity and interplay between molecules and their assemblies are crucial features of biological and chemical systems and understanding its fundamentals is becoming crucial for synthetic supramolecular systems(1, 2). In addition, complex phenomena involving nonlinearity such as competition, self-sorting(3), crosstalk(4), scaffolding(5), templating(6), cooperativity(7), multivalency(8) and ultra-sensitivity(9) often require the aid of mathematical models for detailed analysis and understanding of the crucial molecular interactions involved. In turn, the use of (thermodynamic) computational models has gained popularity to deduce binding mechanisms involved, design experiments, analyze data and determine system constants(10).

The solution to a two-component system with one-to-one binding goes as far back as the Langmuir adsorption model, however more complicated systems consisting of three or more components can only be solved analytically in certain specific cases(11, 12) or after additional assumptions(13). The ternary body problem is mathematically unsolvable without approximation or assumption of the free concentration of one of the components(14). It has therefore gotten more common to develop custom-made mathematical models for specific types of systems based on a combination of analytical and numerical solutions(15, 16). Such equilibrium models have been used for analysis of combinatorial libraries(17), macromolecular reactions within living cells(18), supramolecular copolymerization(19) and the design of synthetic signaling pathways(20). However, the creation of a new model for each unique system takes expertise and a non-trivial amount of time. The widespread integration of these models is thus impeded, even though a trove of information can be extracted from them. Increasing the accessibility of numerical equilibrium models will greatly aid the study of chemical and biological complex molecular systems.

Here we present a framework for equilibrium models as a general approach to generating a custom model for any arbitrary molecular system based on the simple, reversible reactions that constitute that system. The *framework* automatically determines the relations between the species concentrations at equilibrium and combines these with mass balance equations to establish a system of coupled equilibrium expressions that are solved numerically, without the need to rely on assumption or simplifications. In addition, the *framework* facilitates standardized methods for system and parameter analysis. This automated approach severely reduces both the expertise and time required to construct new computational models. The *framework* was designed to cater to both the computational scientist and the experimental (laboratory) scientist, who is possibly less accustomed to programming and modelling techniques.

## Results

We start with a description of our general derivation method. Next, we revisit systems from recent literature to illustrate the simplicity and generality of the *framework* while providing models of the same quality as models created specifically for the cases in question. The first system contains a simple one-to-one interaction and will serve as an introduction to the *framework*. The system complexity is then stepwise increased by revisiting the stabilization of protein-protein interactions(21) and the design of multicomponent homogeneous immunoassays(22). A complete protocol for the *framework* and detailed tutorial descriptions to follow along with each of the cases are available in the supporting information.

### General approach

*The framework* utilizes an automatic model building process, based on a general derivation method in order to streamline the creation of new equilibrium models. A schematic overview of all steps and the workflow are depicted in Figure 1. Reversible chemical reactions are a familiar and natural way to describe interactions in a system and therefore serve as basis for the *framework*. Combined, these reversible reactions form the *system description*, defining the system and all complexes that are formed therein. In a system at equilibrium, each reversible reaction is also in equilibrium and the ratio of species on both sides of each reaction is determined by the equilibrium constant(23). Therefore, the entire system can be described in terms of the equilibrium constants and the free concentrations of species that cannot disassociate further, designated the *components* of the system. The *model builder* determines an expression for the equilibrium concentration of each species in the system based on the equilibrium constant and concentrations of the complexing species. These expressions are reworked until they contain only the *component* concentrations and equilibrium constants.

**Figure 1.**
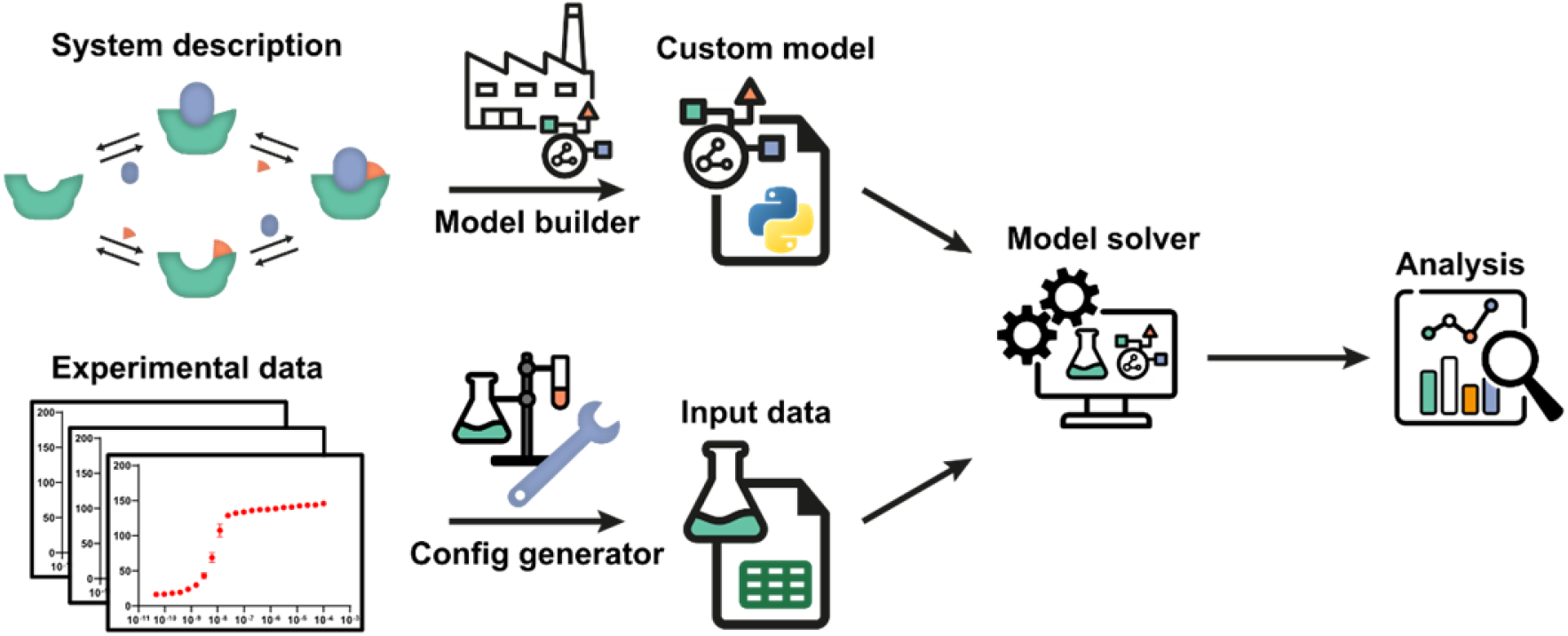
Schematic overview and workflow of the framework. The system description, a set of reversible reactions that describe how all species in the system interact, is translated to a custom model by the model builder. After defining the experimental conditions using the config generator, experimental data can be used by the model solver in order to determine point estimates for any unknown parameters in the custom model. Subsequently, the custom model can be analyzed by the framework. A number of diverse, extendible, and customizable methods are offered.

For each *component*, a mass balance equation is set up. Substituting the expressions of the complexes determined earlier in these mass-balance equations results in a system of *n* equations with the *n* free component concentrations as unknowns. These *equilibrium equations* can then be solved numerically by the *framework*. All steps are performed autonomously based on the entered reversible reactions, generating the desired *custom model*. A comprehensive protocol for each step in the *framework* is in available in SI Appendix, section 1.

Once a *custom model* has been generated, analyses can be performed in the *framework* to gain insight into the molecular system at hand. These analyses include plotting the concentrations of all species in the system as function of *component* concentrations for given equilibrium constants and parameter sensitivity analysis.

The *framework* can also determine unknown parameters by fitting experimental data using the *model solver*. For this, a *data function* is needed that relates *framework* determined concentrations to the measured experimental values. The *data function* is specified in the model builder and is included in the *custom model*. The experimental data values must be a direct function of one or more species concentrations(10, 23). Experiments where this is the case often include titrations and we will use the related terminology in the given examples. Experimental conditions in which the data was obtained are defined with help of the *config generator*. The *framework* protocol contains instructions and the expected data format. The *model solver* will then calculate the equilibrium concentrations of all species based on the specified total concentrations, and initial guesses for the unknown parameters, for each datapoint. Parameter values are iteratively adjusted to best fit the entire experimental dataset using a numerical least-squares optimization.

The *framework* also offers analyses to assess the quality of the determined parameter estimates, including model prediction plots, mean-squared-error landscapes and confidence intervals determined using the bootstrap method. A complete overview of all analyses is available in SI Appendix, section 2. In the next sections we will apply the *framework* to increasingly complex systems by first constructing a *custom model* and preparing the experimental data, comparing the parameter estimates determined to previous findings and finally highlighting some of the analysis methods the *framework* can provide.

### Binding between the 14-3-3 protein and a partner protein (3 species, fluorescence anisotropy data)

The first system consists of the one-to-one interaction between the protein 14-3-3 and one of its protein targets TASK3, a potassium ion channel. 14-3-3 proteins are a family of highly preserved scaffold proteins that interact with hundreds of distinct protein partners(24). These hub proteins are involved in processes such as cell signaling, protein trafficking, cell cycle progression and apoptosis(25, 26). Because of this, 14-3-3 proteins are closely involved with a number of human diseases and have proven to be interesting drug targets(26). Here, we will determine the dissociation constant for the interaction between 14-3-3 and the TASK3 protein based on fluorescent anisotropy data (Dataset S1)(27). This system serves as an introduction to our *framework* approach, as such one-to-one interactions can also be solved analytically(28).

The system consists of only a single reversible reaction and the *system description* is depicted in Figure 2A. The next step in modelling this system is creating the *custom model*. Each reaction is entered as ‘complexing species = complex; dissociation constant’ (see Figure 2A). In addition to the protocol, the SI contains an overview with the exact input values for each step to model the systems discussed here, which can be used as a tutorial reference (SI Appendix, section 3). From the system description, the equilibrium equations in Figure 2B are generated. Note how the complex concentrations are substituted so the final equations only consist of *component* concentrations and equilibrium constants.

**Figure 2:**
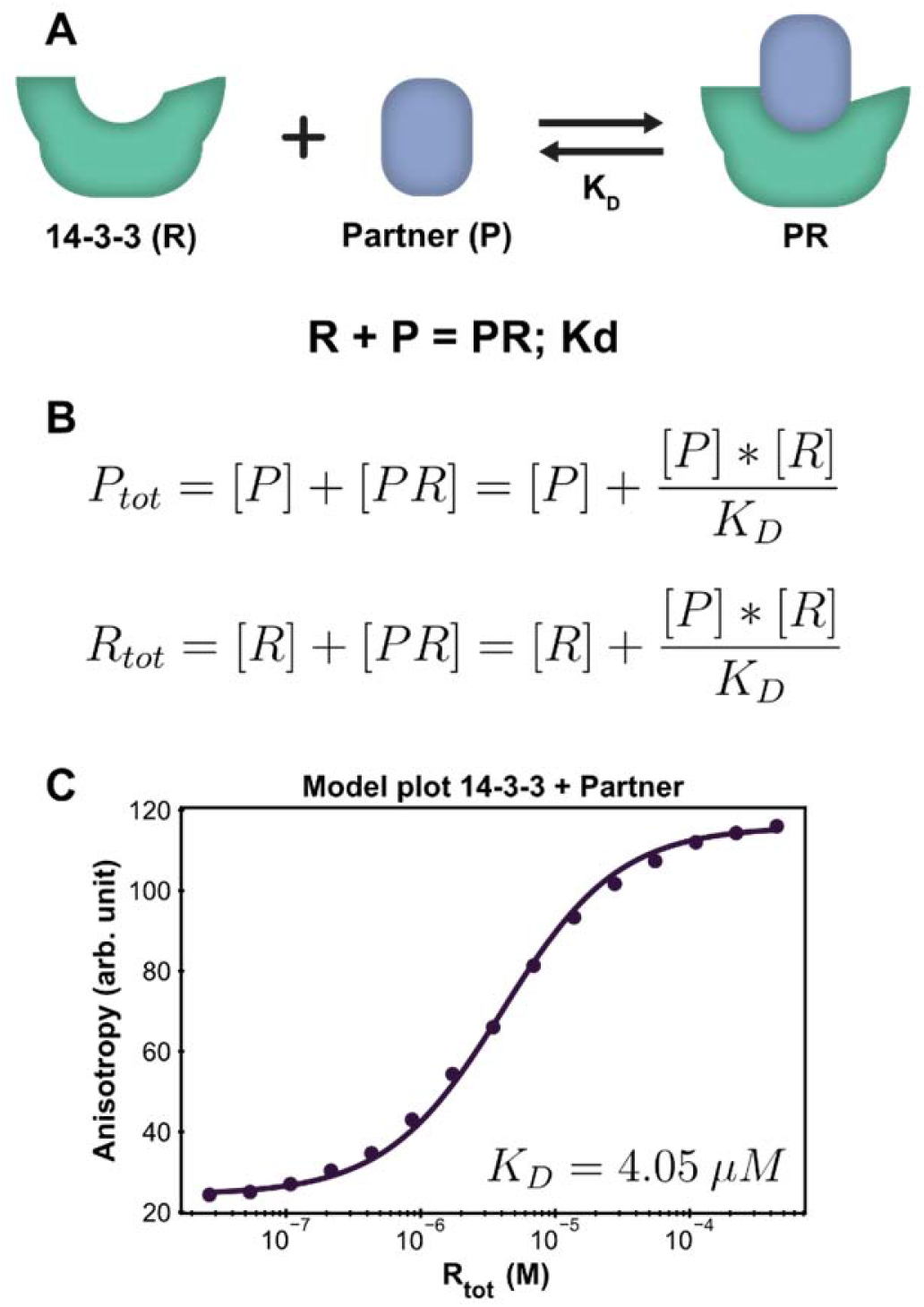
Application of the framework on the introductory system. **A:** System description, each 14-3-3 monomer (R) can complex with one partner (P) molecule (TASK3). The model builder input corresponding to this reversible reaction is displayed beneath the cartoon. **B:** The equilibrium equations determined for this system, before being rewritten for numerical efficiency. **C**: Fluorescence anisotropy data collected for this system(21) (dots are the averages of the experimental technical repeats) in the presence of 10 nM partner (labeled component) together with the model fit (line).

After setting up the *custom model* and preparing the experimental data, parameter fitting and additional analysis is performed from the *framework* main script, which offers a simplified interface to the underlying functions. It is also possible to directly execute the functions in the *framework* for advanced customization if desired. For this simple introductory system, only the model plot analysis will be executed. The *framework* will determine point estimates for the fit parameters before performing any chosen analyses. It is important to consider that the optimal estimates are determined within the constraints of the *custom model* and the *framework* does not judge the correctness of the *system description*. In order to validate the proposed *system description*, we inspect both the parameter values and the model plot. The calculated estimate for the *K_D_* is 4.05 *μ*M, which is in line with previous findings(29). The chosen analysis visualizes all the experimental data and the model prediction for each titrate (14-3-3) concentration using the determined point estimate (Figure 2C). Inspecting the graph shows that the model prediction accurately describes the data with the determined parameter value.

### Protein-protein interaction stabilization (6 species, 2D fluorescence anisotropy data)

For the second case, the previous system will be extended with a third molecule that interacts with the binary protein-protein interactions system to illustrate the capabilities of the framework in a more complex ternary body problem setting. Such 14-3-3 ‘molecular glues’ are an emerging and versatile strategy in drug development(30, 31). At its core, the interaction between two proteins is selectively stabilized by way of a third, low molecular weight compound(32). Besides the affinity of the stabilizer for the protein target, the measure of cooperativity induced by the stabilizer is also critical optimization parameter. The extended system definition can be seen in Figure 3A. While it is still possible for 14-3-3 (R) to directly bind to the partner protein (P), an additional path is now available where a stabilizer molecule binds first. As a result, the affinity for the partner is greatly increased. Because of the thermodynamic cycle in this system, it can be described entirely using the individual binding affinities (K*_D,I_*, *K_D,II_*) and a cooperativity factor (*α*)(21). In this system, the stabilizer molecule (S) does not interact significantly with the partner protein by itself. The experimental data for this case consists of 2D fluorescence anisotropy titrations (Figure 3B, Dataset S2)(21). The goal is to determine point estimates for the cooperativity parameter and the binding affinity of the stabilizer for the combined dataset and analyze the quality of these estimates.

**Figure 3:**
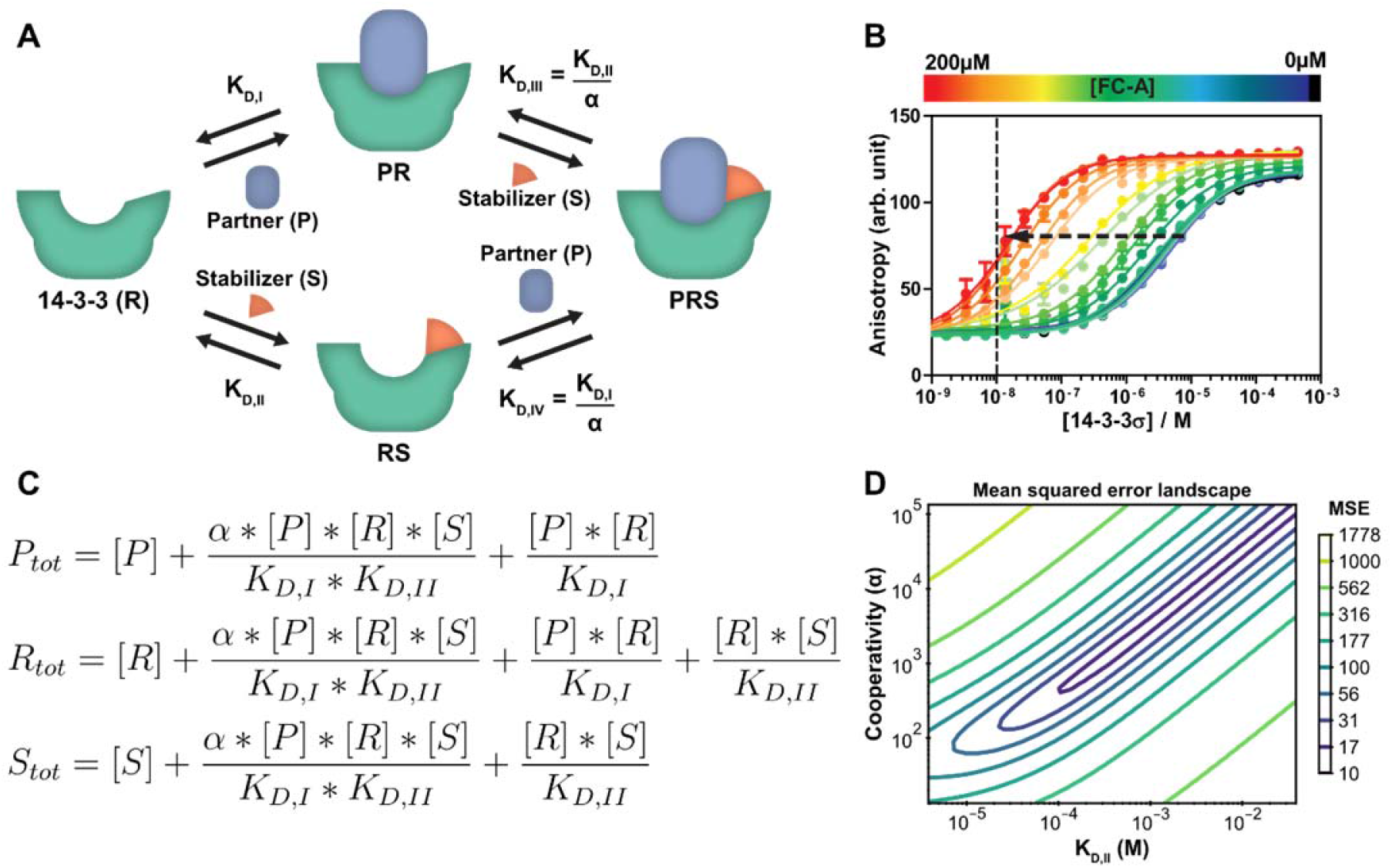
Protein-protein interaction stabilization case. **A:**System description(21): the adapter protein 14-3-3 (R) binds to its partner TASK3 (P) with dissociation constant *K_D,I_*. In the presence of fusicoccin-A (S), the interaction is stabilized by a factor *α*, reducing the apparent affinity. Either the partner or stabilizer can bind first to form the ternary complex. **B:** Fluorescent polarization data in the presence of varying concentrations of fusicoccin-A and a partner (labeled component) concentration of 10 nM (line), previously published(21) for the model created specifically for this system. Note the decrease in EC_50_ value with the addition of the stabilizer (arrow). **C:** Equilibrium equations automatically determined by the framework for this system. **D:** Error-landscape plot centered on the determined estimates. The contours show that there is a valley of parameter combinations that result in a relatively low mean squared error (MSE).

A *custom model* is created for the *system description* displayed in Figure 3A, resulting in the *equilibrium equations* of Figure 3C. This three *component* system results in three distinct equations. The step-by-step guide for this system can be found in SI Appendix, section 4. For this molecular system, one of the parameters is already known (*K_D,I_*, 4.05 *μ*M) and as such entered and kept constant while the *model solver* determines point estimates for the *α* and *K_D, II_*, parameters, resulting in values of 1.34 × 10^3^ and 0.389 mM respectively. These are in good agreement with the previously determined values of 1 × 10^3^ and 0.3 mM(21). Thus, even equilibria with more complex *system description* and experimental data can be modeled correctly using the *framework*.

Next, we visualize the mean squared error landscape around the determined point estimates in order to gain more insight into the parameters and the determined point estimates (Figure 3D). The landscape gives a sense of the interdependence of the two parameters and the influence of each by displaying contour lines. A valley of parameter combinations results in a relatively low mean squared error. The *K_D,II_* and *α* values display positive correlation, increasing both values (weaker initial binding, stronger cooperativity) still results in a relatively good prediction of the input data. Nevertheless, starting the solver from several combinations found at the valley of the landscape as initial guess values results in the same point estimates, indicating that there is a small preference for the determined estimates over the other possible combinations. A sharp rise in error can be observed when one of the parameters is fixed and the other is varied. This indicates that the ratio of these parameters is important for accurate model prediction and can thus be determined with high confidence.

While a point estimate is the best single-value approximation of a parameter, a confidence interval can be effective to get a sense for the certainty (or spread) of the reported estimate. The *framework* can determine this interval using a nonparametric, bias corrected bootstrap approach(33–35). As an example, the confidence interval analysis is performed with 2000 repeats in order to get an appropriate sample size and a 95% confidence interval (SI Appendix, Figure S6). Additional information on the bootstrap method is available in SI Appendix, section 5. The confidence interval (lower; median; upper) for the *K_D,II_*, (0.189 mM; 0.392 mM; 2.19 mM) and the *α* (0.687 × 10^3^; 1.34 × 10^3^; 7.35 × 10^3^) show an order of magnitude difference between the limits of the confidence interval. This broad range corresponds with the stretched valley in Figure 3D and is important to consider when drawing conclusions from parameter estimates.

### Ratiometric Plug-and-Play Immunodiagnostics platform (10 species, luminescence data)

The last exemplary molecular system will demonstrate the *framework’s* capacity for extended systems and displays some of the more advanced features. In addition, we will demonstrate how the *custom model* can guide system engineering and influence experimental design. The recently developed Ratiometric Plug-and-Play Immunodiagnostics (RAPPID) platform will serve as context for this discussion(22). RAPPID facilitates the development of ratiometric bioluminescent immunoassays for a wide range of biomolecular targets. The platform combines the use of antibodies with split NanoLuc luciferases(36, 37) to detect the formation of sandwich immunocomplexes in solution. A schematic overview of the general system is depicted in Figure 4A. The system consists of two types of antibodies (A, B) and an analyte (T). Each antibody binds to a different epitope (with affinities *K_D,A_* and *K_D,B_*) which allows for the formation of the ABTi(nactive) sandwich. The two parts of the spilt NanoLuc have a designed affinity of *K_D,N_*. The increased effective molarity (EM) within the ternary complex promotes the subsequent complementation into the ABTa(ctive) NanoLuc complex, which then emits light. Statistical factors are present in this model to derive the dissociation constants *K_D,A_* and *K_D,B_* as there are two possible binding sites between each bivalent antibody and the monomeric target(22, 23). At higher analyte:antibody ratios, the formation of analyte-antibody-analyte complexes becomes predominant over the formation of the functional ternary complex, giving rise to the so-called hook effect(38).

**Figure 4:**
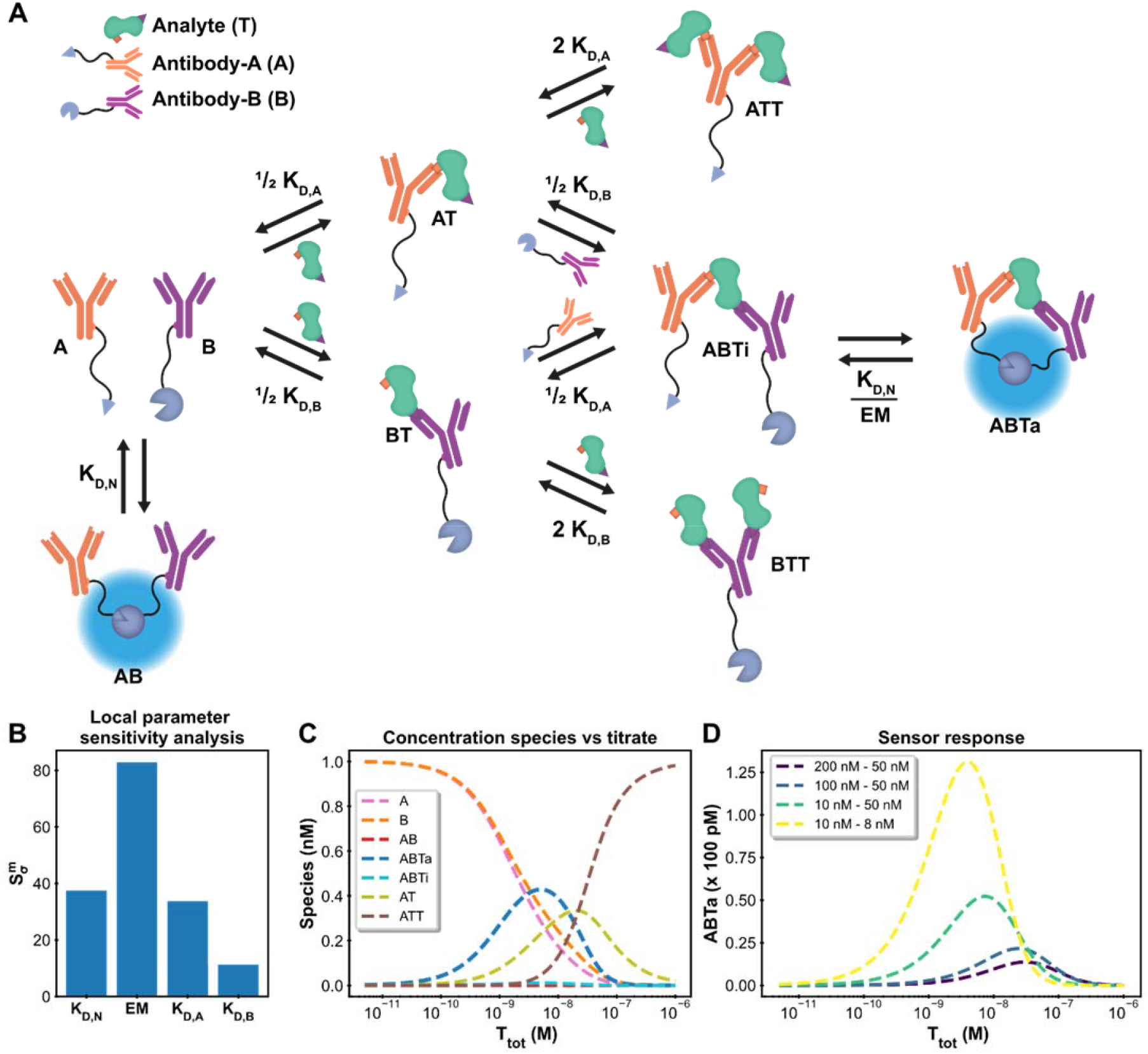
Ratiometric Plug-and-Play Immunodiagnostics (RAPPID) case. **A:** System description(22). The antibodies A and B are both conjugated to a part of split NanoLuc luciferase, which emit light upon complex formation. The antibodies are specific for a certain analyte but target different epitopes. Upon antibody-analyte sandwich formation, the effective molarity for the split luciferases increases, resulting in increased complex formation and consequently increased signal. **B:** Local parameter sensitivity analysis for the system using cardiac troponin I data (and corresponding concentrations) and a perturbation of +50%. The reference values are the point estimates determined by the framework. **C:** Concentration plot generated with framework displaying how each concentration changes as the total titrate concentration increases. This analysis is performed without any experimental data with the fixed parameter values *K_D,N_*: 2.5 μM, *K_D,A_*: 10 nM *K_D,B_*: 15 nM, *EM*: 100 μM, [*A*]_*tot*_: 1 nM, [*B*]_*tot*_: 1 nM. The species BT and BTT have been omitted for clarity. **D:** Simulation of shift in active ternary complex (ABTa) concentration for different combinations of antibody affinities and an *EM* value of 10 μM. Both antibodies have a total concentration of 1 nM.

We start by comparing the *framework custom model* with the model for intensiometric detection of cardiac troponin I, presented in the RAPPID manuscript. Experimental data (Dataset S3) and step-by-step guide (SI Appendix, section 6) are also available. The *system description* is displayed in Figure 4A. A total of eight reversible reactions describe this system. The measured data (SI Appendix, Figure S8) is proportional to the concentration of formed active ternary complex (ABTa) multiplied by an unknown scaling factor, representing the enzymatic activity of the luciferase. This data-concentration relationship is not available as a default data function in the *model builder*. However, the *framework* is not limited to these predefined functions, and it is possible to easily specify the relationship. In the model builder we define the custom function: *ABTa * Scaling*. Instructions on custom data functions are available in SI Appendix, section 7. Determining point estimates for the parameters *K_D,A_*, *K_D,B_* and *Scaling* with fixed values for *K_D,N_* and *EM* results in values of 533 nM, 15.3 nM and 5.53 × 10^17^ RLU / M respectively, which are identical to the values determined in the original paper(22). As such, using the *framework* even complicated multi-component systems are easily modelled and with equal accuracy as specifically designed models.

Usually, not all parameters contribute equally to the final measurement or result of a model. The influence of each parameter can be determined using local parameter sensitivity analysis(39). This quantifies the change in a given function (designated M) based on a percentage change in parameter value. The default M-function measures the change in the sum of squared errors between the model prediction and the experimental data. Parameters with high sensitivity greatly affect the final model prediction. Figure 4B displays the sensitivity for the parameters in the cardiac troponin I model after a 50% increase in the parameter value. The sensitivity of *K_D,A_* is larger than *K_D,B_*. Because the binding affinity of the A-antibody is significantly weaker than that of the B-antibody it serves as bottleneck in the formation of the active complex. In addition, both the *K_D,N_* and the *EM* parameters, which are fixed parameters, have greater sensitivity than the fitted parameters (*K_D,A_, K_D,B_*). An error in the fixed values of sensitive parameters will greatly affect the determined estimates for the other parameters. Obtaining accurate values for these parameters is therefore critical and experimental setups can be adjusted to suit these criteria. Another example which measures the sensitivity for the maximum amount of active complex formation is given in SI Appendix, section 8.

For the design of new molecular sensors, it is important to engineer the detection regime over a large range of possible analyte concentrations. Within the RAPPID system, the specific antibodies and the concentrations of the sensor components can be relatively easily adjusted in order to tune the sensor. While it is possible to gather new experimental data, it is more (cost-)efficient to use the *framework*. We use the concentrations analysis for this purpose. In this analysis, the concentrations of all species in the system over a range of titrate values is visualized, (Figure 4C). The final result for a number of antibody affinities can be seen in Figure 4D. The peak of the graph, where the greatest increase in luminescence signal is observed, is dictated mostly by the antibody affinity. Greater affinities also increase the total signal as can be observed from the figure. The maximum complex formation can be tuned by changing the total antibody concentration(22). The analysis allows the selection of antibodies that are most suited for the intended analyte concentrations at a fraction of the time or costs necessary to perform the experimental measurements.

When building a *custom model* by hand for a more elaborate multicomponent system it is not uncommon to exclude certain (higher-order) complexes that are assumed to form in negligible amounts as manually deriving the *equilibrium equations* for these larger systems is labor intensive and error prone. Using the *framework* approach it is possible to extend any multicomponent system within minutes which allows for proper validation of such assumptions. An extended system description the RAPPID system was modeled in SI Appendix, section 9. This extended *custom model* revealed that several additional four-component complexes are formed in significant amounts at higher analyst concentrations and should potentially not be ignored. Fitting parameters to the extended *custom model* results in the point estimates *K_D,A_* = 46.7 nM, *K_D,B_* = 9.71 nM and *Scaling* = 3.75 × 10^16^ RLU / M. These values have up to an order of magnitude difference compared to the previous estimates. This shows that the *framework* is not only capable of modelling complex systems, but also facilitates simple and fast validation of existing models.

## Discussion

Equilibrium models can provide a wealth of information about molecular systems such as of the medicinal chemistry, supramolecular, or biochemical type. However, their creation, including the derivation of equilibrium equations, takes expertise and a non-trivial amount of time. The *framework* approach is capable of generating equilibrium models using an automatic derivation process for arbitrary molecular systems, defined by reversible reactions. A side-by-side comparison with three systems of increasing complexity demonstrates the scope and accuracy of the *framework*. The parameter point-estimates determined by the *framework* closely matched or were identical to the values determined by the original models created solely for each of the systems in question. The *framework* can also be used to gain additional insight into the quality of the determined estimates and the interrelationship of parameters in the model. Furthermore, building and analyzing a computational equilibrium model using the *framework* can optimize experimental design and reduce the need for multiple experiments, even in situations without any prior experimental data.

The *framework* features an easy-to-use design that does not require a programming or mathematics background. Advanced users will find that the general structure and use of the Python programming language allows for straightforward extension and customization towards specific usage scenarios. The *framework* thus facilitates the simple and fast creation of effective computational equilibrium models in order to unravel, understand and delineate a broad range of molecular systems.

## Methods

The *framework* is constructed entirely in Python and is freely available from **DOI:**10.5281/zenodo.5531622. A complete overview of all required dependencies can be found in SI Appendix, section 1. The solver section uses an object-oriented approach and supports modification of parts for specific use cases while the analysis section uses separate functions to facilitate addition of custom analysis functions with automatic integration into the rest of the *framework*. Users familiar with Python can easily add specific analyses appropriate for their research as needed (SI Appendix, section 10). Parameters are determined using the iterative scipy.optimize.least_squares solver with the ‘Trust Region Reflective’ algorithm(40). In addition, by default a log transform(41, 42) is applied to all parameters to support large order-of-magnitude differences between parameters and to increase the solving speed. The error function is defined as the mean squared error between the experimental values (averaging technical repeats) and the data-function prediction for each titrate concentration. Multiple methods are implemented in order to solve the equilibrium equations (model.system_equations) for a specific titrate concentration. The solver will first attempt to use the faster but less robust scipy.optimize.root method before escalating to a least-squares approach in case no physical equilibrium can be determined. Based on the solution, all species concentrations are updated to the new equilibrium in the state object (model.update_state) and the data function (model.data_function) converts species concentrations to an experimental data value prediction which is used in the error function. Stress tests have been performed to test the limits of the *framework* (SI Appendix, section 11). Systems of up to a hundred unique species were tested and produced solutions in a reasonable time span.

## Data availability

Al study data is included with the article.

## Acknowledgments

This research was funded by the Netherlands Organization for Scientific Research (NWO) through Gravity program 024.001.035, VICI grant 016.150.366 and the European Union’s Horizon 2020 research and innovation program under the Marie Skłodowska-Curie grant agreement No 844872 (TW). We thank Marloes Pennings and Kilian Cozijnsen for their feedback as beta-users and Maarten Merkx and Bas Rosier for the fruitful discussions regarding the RAPPID model.

