## Supporting Information for "A general framework for straightforward model construction of multi-component thermodynamic equilibrium systems"

**Contents:**

Supplementary sections 1 to 11
Figures S1 to S12
SI References

**Other supplementary materials for this manuscript are the framework and data sets. These are freely available from:**

DOI: 10.5281/zenodo.5531622

**Contents**

1. Framework for equilibrium models, protocol ... 4
    - Introduction ... 4
        - Required expertise ... 4
        - Limitations ... 4
    - Materials ... 5
        - Experimental data ... 5
        - Required hardware ... 5
        - Required software ... 5
        - Optional software ... 5
    - Procedure ... 5
        - System definition and experimental data (2 – 10 min) ... 5
        - Loading the model builder (10 s – 2 min) ... 6
        - Creating a custom model (4 - 15 min) ... 6
        - Preparing experimental data and config ( – ) ... 8
        - Determining point estimates and analysis (2 – 5 min) ... 9
    - Model builder rules ... 10
2. Analysis method overview ... 12
3. Case I: Binding between 14-3-3 protein and partner protein, step-by-step guide ... 14
    - System description (step 1) ... 14
    - Model builder (steps 2-8) ... 14
    - Experimental data and config file (steps 9-11) ... 14
    - Main script (steps 12-21) ... 15
    - Check model (step 22) ... 16
4. Case II: Protein-protein interaction (PPI) stabilization, step-by-step guide ... 17
    - System description (step 1) ... 17
    - Model builder (steps 2-8) ... 17
    - Experimental data and config file (steps 9-11) ... 18
    - Main (steps 12-21) ... 19
    - Check model (step 22) ... 20
5. Bootstrap ... 21
6. Case III: Ratiometric Plug-and-Play Immunodiagnostics platform (RAPPID), step-by-step guide 22
    - System description ... 22
    - Model builder (steps 2-8) ... 22
    - Experimental data and config file (steps 9-11) ... 23
    - Main (steps 12-21) ... 24

### 1. Framework for equilibrium models, protocol

#### Introduction

The *framework* supports parameter estimation and analysis of thermodynamic equilibrium systems without prior modelling experience (Figure S1). The process starts with the definition of the possible specie interactions within the system of interest (referred to as the *system description*) and in case parameters are fitted, experimental data for the system. The *system description* is specified using a number of reversible chemical reactions. The *model builder* is able to automatically determine mass-balance and *equilibrium equations* from such reversible reactions and creates a *custom model* which can be loaded into the *framework* for analysis. Unknown parameters are determined using an iterative least-squares optimization based on the experimental data entered. It is important that the measured values within the experimental data are a function of one or more specie concentrations in the system. The *framework* also offers analysis functions in order to e.g. determine relationships between parameters, sensitivity of individual parameters or confidence intervals based on a bootstrap approach.

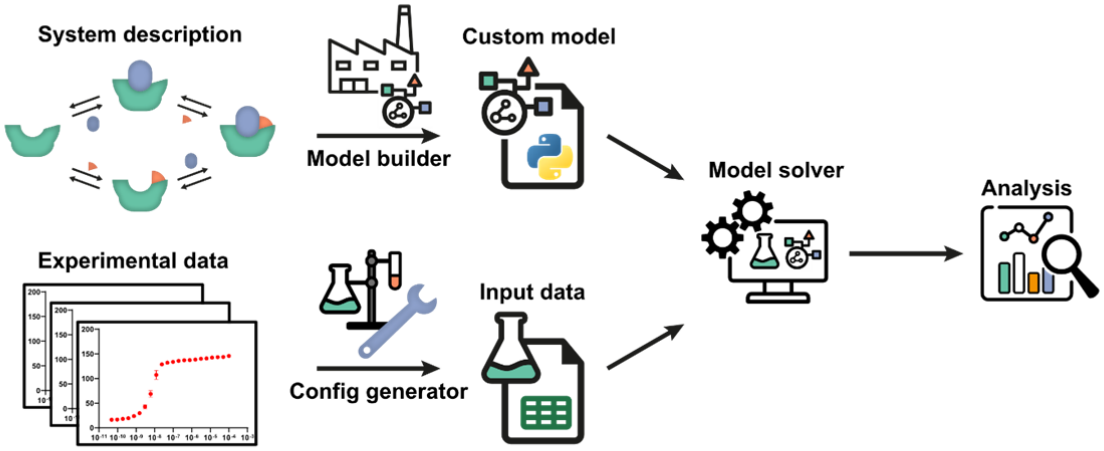

Figure S1: **Schematic overview and workflow of the framework as depicted in the main text.** The system description, a set of reversible reactions that describe how all species in the system interact, is translated to a custom model by the model builder. After defining the experimental conditions using the config generator, experimental data can be used by the model solver in order to determine point estimates for any unknown parameters in the custom model. Subsequently, the custom model can be analyzed by the framework**.** Several diverse, extendible, and customizable methods are offered.

#### Required expertise

Most of this protocol can be implemented without any prior knowledge of model building or the Python programming language. An introductory level of familiarity with the Python syntax and standard types (e.g. list, string, int) will help prevent common errors when entering parameter / option values. The *framework* is set up in such a way that more advanced Python users have the ability to customize the execution of some functions or add new functionalities with easy integration into the *framework*. This protocol focuses on the basic procedure. Adding custom analyses is discussed in section 10. A standard desktop computer is ample to run the framework and working knowledge of the installed operating system is sufficient.

#### Limitations

Systems consisting of non-reversible reactions cannot be created using the *model builder*. The *framework* assumes equilibrium conditions and determines the best fit for the given *custom model*. If the model is incorrect, the parameter estimations will also be incorrect. Always critically review the determined estimates in order to judge whether they are plausible based on previous research.

**CRITICAL** Parameters that significantly differ from expected values are substantial grounds to review the proposed *system description*.

### Materials

#### Experimental data

Experimentally obtained values should be a function of one or more specie concentrations within the system. Titration experiments are especially supported by the framework.

#### Required hardware

- A standard desktop computer or laptop will be sufficient for most functionalities. In order to determine confidence intervals for larger datasets, a desktop computer is recommended due to the increased computational complexity.
- A hard drive with at least 10 GB of free storage if Anaconda has not been previously installed.
- A working internet connection is required during installation.

#### Required software

A distribution version of the framework, available from **DOI:** 10.5281/zenodo.5303678.

In order to use the *framework*, Python version 3.8 or newer, and a number of common Python packages are required. **CRITICAL** For users with limited experience with the Python environment the authors highly recommend the Anaconda package (https://www.anaconda.com/products/individual). The Anaconda package includes all required software for the model framework in a single installation. Help on installing the anaconda package can be found on https://docs.anaconda.com/anaconda/install/. For users who prefer not to install the Anaconda package, the following software should be installed by other means:

- Python, version 3.8 or above.
- NumPy, version 1.20.1 or above, provides array processing and is a fundamental package for scientific work with Python.
- SciPy, version 1.6.2 or above, is a scientific computing package which provides the solvers to determine equilibrium conditions.
- Pandas, version 1.2.4 or above, is a high-performance data structure and data analysis tool.
- Matplotlib, version 3.3.4, provides MATLAB-style high-quality figures in Python.
- SymPy, version 1.8, is required in order to construct models using the 'model builder' tool.

#### Optional software

i. The current version (1.x) of the *framework* does not offer a graphical user interface. It is therefore recommended to install an integrated development environment (IDE) in order to assists in editing input parameters and in analysis. If the Anaconda package was installed, Spyder (https://www.spyder-ide.org/) will be installed and available for this purpose. Other potential options include, but are not limited to, Eclipse (https://www.eclipse.org/) and PyCharm (https://www.jetbrains.com/pycharm/).

### Procedure

The entire procedure can be divided into five general steps. I: defining the *system description*, II: creating a *custom model*, III: preparing experimental data, IV: determining point estimates and V: performing analysis. Each step will be discussed in turn.

#### System definition and experimental data (2 – 10 min)

*The* system description *defines how the species within the system can interact and can form complexes. It forms the basis for the model that will be constructed.*

1. **Define the system** – By using reversible chemical reactions, define all interactions in the system. It is recommended to either draw the system or construct a computational image. Note reactants, products and reaction rates. **CRITICAL** As is customary in biochemistry, constants are expected to be defined as dissociation constants ($K_D$). An example definition of a relatively small system can be seen in Figure S2.

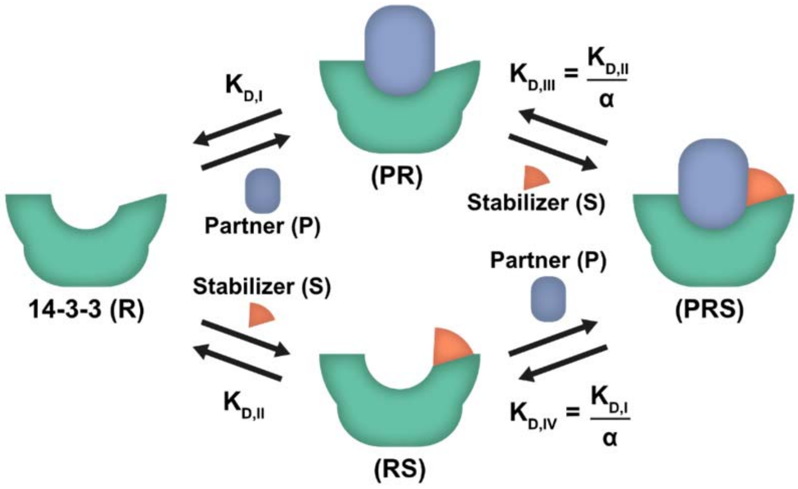

Figure S2 **Example system description.** Note that all components have been given names (in brackets) and that for each reversible reaction an equilibrium rate constant has been defined.

### Loading the model builder (10 s – 2 min)

2. **Open the 'model builder.py' script** – For users unfamiliar with Python, the Spyder IDE has been installed during installation of the Anaconda package which can be used for this purpose. The procedure will focus on specific actions required in the Spyder environment as it is expected that more advanced users will be able to locate the corresponding actions in their editor of choice.

   **Optional: Using Spyder to open scripts**

   *i.* Open the Spyder IDE. Windows users can open the start window and type 'Spyder' in the search bar. Click the displayed Spyder icon. Alternatively, the anaconda manager can be started from which Spyder can be started.

   *ii.* In the top left, press the folder icon, alternatively go to File → Open…

   *iii.* Browse to the location where the *framework* files are extracted.

   *iv.* In the main folder, open the "model builder.py" script by double-clicking it.

### Creating a custom model (4 - 15 min)

*The* custom model *is the version of the system description that can be understood and loaded by the* framework*. It also contains the data function which maps the system's specie concentrations to measured values. Systems that interact in the same way (e.g. multiple potential drug compounds) only require the creation of a single* custom model*.*

3. **Change the name for the new model** – A *custom model* file defines how the species in the system interact. A model file only needs to be created once for each distinct system. **CRITICAL** Systems in which components interact in the same way can use the same model file (e.g. for testing different potential drug compounds that act on the same target). In the previously opened 'model builder.py' script change the name of the new model to one of your choice. Remember to leave the quotation marks around the name, see **Intermezzo: Python types**

*Intermezzo: Python types*

While a very basic knowledge of Python is expected in this procedure, here is a refresher on how some basic types of data are entered in Python.

**Text** (a sequences of characters) is entered as a string. In order to tell Python something is a string, encase it in single or double quotes.

e.g. "My model file name"

**Whole numbers** are saved as integers. These can be entered by just typing the number.

e.g. 42

**Decimals** are saved as floating-point numbers (floats). Two ways to enter decimals are either by using the '.' notation or by using scientific notation.

e.g. 0.001, 15E-6.

**Multi-line text** is similar to text but spans more than a single line. In order to enter text that continues over multiple lines place three double-quotes """…""" around the text instead of single quotes.

**Pairs of values** are saved in a **dictionary**. Within the *framework* these are used to define options or assign values to parameters. Within this context a dictionary looks like this: {parameter: value}, each pair of values is separated by a comma, e.g. {'Alpha': 50, 'KD2':5E-6} (remember the quotes around text).

4. **Enter the system description equations** – In order to enter the equations which will define the model, several rules need to be followed for correct definition. For the complete overview, see **Model builder rules**. First-time users are recommended to read these specifics before continuing. For each reversible reaction in the system defined in step 1, one line will be added. Leave the triple quotes around the equations. For the example system in Figure S2, the following lines would be entered (the order in which the lines are entered does not change the model definition):

```
system_description = ("""
R + P = PR; Kd1
PR + S = PRS; Kd2 / Alpha
R + S = RS; Kd2
RS + P = PRS; Kd1/ Alpha
""")
```

**CRITICAL** Note that the input is case sensitive and that each reversible reaction in the defined system is entered on a new line.

5. **Choose data function** – Choose the data function which suits the experimental data. The data function determines the experimental value corresponding to a specific combination of specie concentrations and parameter values. This function is used to fit the experimental data. A list of available options is given in the model builder script. Change the data function to the applicable one, leaving the quotes. In case no description fits the experimental setup, see **Custom data functions** for details on entering a custom data function.
6. **Define the labelled specie** – From the species entered in step 4, enter which component is labelled or tracked in the experiment by changing the 'labelled' value. Not required when the 'custom' data function is chosen.
7. **Run the python script** – for Spyder users, this can be done by pressing the green arrow in the top-left or by pressing F5.
8. **Confirm the results and copy custom model** – The model builder will print information about the generated model to the console (lower right in the default Spyder layout). **CRITICAL** Carefully inspect the information, any unexpected values might be caused by errors in the system description equations. Make a note of the reported components, these will be needed in a later

step. If all the information is correct, open the file explorer and navigate to the model framework location. Move the created file from the 'output' folder to the 'model' folder.

Preparing experimental data and config ( – )

*When there are parameters within the system for which no known value is available, the* framework *can fit these values based on experimental data. The framework requires a structured format in order to correctly analyze the data. Alternatively, the* custom model *can be analyzed without any data based on estimates / known values for all parameters. In both cases, a config file is used to define the total concentrations of the components in the system.*

9. **Optional: gather measurement data** – Multiple experiment types are supported but it is critical that the measured value is a function of one or more concentrations within the system. An example experiment type is fluorescent polarization. In fluorescent polarization measurements, the output signal is proportional to the concentration of labelled specie that is bound to other species in the system.
10. **Optional: Preparing experimental data** – The *framework* requires a structured input format in order to correctly fit the data. This description will focus on Microsoft Excel, but the OpenDocument Spreadsheet format is also supported. The first column of the file should contain titrate concentrations **CRITICAL** in molar. The following columns contain measurement data. The first row contains headers which separate multiple experiments. See the *'example_input.xlsx'* example file. If a program such as GraphPad was used, it is usually possible to copy all the data without any additional modifications.
    i. **Copy the data to excel** – Open a new excel file and use the first sheet. Follow the guidelines given above. Each group of technical repeats should contain the name of that group in the first (left-most) column of that group, e.g. for the triplo measurement E-G, make sure the name is in E1. Should the copy source change these header cells into a single cell (as is the case for e.g. GraphPad) than this will not lead to issues. Missing values can be entered by leaving a field empty. **CRITICAL** Make sure each group of technical repeats has a name, otherwise they are seen as one large group of technical repeats. Save the file.
    ii. **Open '*generate config.py*' script** – For help on opening files in Spyder, refer to step 3.
    iii. **Change input location** – Change the path given after 'input_file' to the saved file, e.g. *"input/my_excel.xlsx".* It is also possible to input direct paths for more experienced users.
    iv. **Run the script** – For Spyder users, use the icon in the top-left or press F5.
    v. **Move the created config** – A 'config.ini' file has been created in the './output' folder for the supplied experimental data. This file is used to define the experimental conditions for each titration. Move the file to the './input' folder for the system.
11. **Define concentrations in config file** – The total concentrations of all components of the system (see the *model builder* steps) need to be defined in order to analyze the system. The titrate concentration is defined within the excel file, the other species will need to be defined using the *'config.ini'* file. *Note: When analyzing a custom model without experimental data, the config file './ _example files/config.ini' can be used. Copy this file to the './input' folder for this system. Place all concentrations under the default header in this case.* Open the file using either Spyder or a text editor such as notepad. A description of how to enter values is given at the top of the file. Values added in the DEFAULT section will be added to all experiments. These will be overwritten by values in experiment specific sections. For the example system given earlier, the partner (P) total ('_tot') concentration is equal for all conditions, but the total stabilizer (S) concentration

differs per condition. An example config file is shown below **CRITICAL** note that the section names correspond to the names supplied in the headers of the Excel file with the experimental data and all text is case sensitive.

```
# Config.ini - protocol example

[DEFAULT]
P_tot = 10E-9
[PPI_1]
S_tot = 1E-9
[PPI_2]
S_tot = 4E-9
[PPI_3]
S_tot = 8E-9
```

The above lines will define the total concentration of P to be 10 nM for all conditions. The total concentration of S is condition specific and is set in the condition specific sections. **CRITICAL** Save the file, the *framework* will raise an exception in case not all required values for solution determination are set.

Determining point estimates and analysis (2 – 5 min)

Framework *combines the* custom model *and* experimental data *in order to determine point estimates for the fit parameters. The* framework *can then be used to perform additional analysis.*

12. **Open the model framework 'main.py' script** – For help on opening files in Spyder, refer to step 3. Upon first time opening this file, example values are present in all fields. Take careful note of the formatting if you are unfamiliar with Python. This script is a convenience interface to the underlying *framework* code.
13. **Enter the model file to use** – Determine which *custom model* to use. No extension ('.py') is required. The model should be located in the './model' folder of the *framework*.
14. **Define the input folder to use** – The input folder contains the 'config.ini' and, when data is fitted, the experimental data file. The default input folder is 'input/'. In the situation where you would like to keep separate folders for different systems / experiments, the input folder to use can be changed here.
15. **Enter the setup type** – See the description in the file for the available modes. Each type is tailored to specific modelling goals.
16. **Enter the titrate parameter** – This is the parameter which corresponds to the 'concentration' value in the input data (the titrate). **CRITICAL** Make sure to enter the parameter which corresponds to the *total* concentration, not the *free* concentration, by adding '_tot' to the end of the name, e.g. 'R' becomes 'R_tot'.
17. **Determine parameters to fit with initial guess** – Fit parameters are the constants in the model for which the framework will determine point-estimates. Note that the parameters should be entered in a dictionary format (see **Intermezzo: Python types**). Each parameter is paired with an initial guess for that parameter. These are the starting points from which the model will determine the point estimates. **CRITICAL** While these estimates do not need to be accurate, there is a chance that multiple minima exists (possible 'false' solutions) and therefore it is recommended to base the initial guess on previous research if available. Alternatively, the 'range_solver' analysis can be used to start solving from a range of possible values in order to assess if they converge to the same point. For the example system we want to fit the parameters 'Alpha' and 'Kd2' resulting in the following input: `{'Kd2':1E-5, 'Alpha':30}`. If no parameters should be fitted, enter an empty dictionary: `{ }`.

18. **Enter known parameter values** – On the next line, using the same paired format, enter any remaining parameters for which the value is known or who's value should remain fixed. If there are no remaining parameters, enter an empty dictionary: `{ }`. For the example system 'Kd1' is known to be 4.05 µM, thus: `{'Kd1': 4.05E-6}`.
19. **Determine analysis to perform** – The analysis list determines which analysis / plot functions will be executed after determining point estimates for the fit parameters (if any). Performing analysis is optional. When no analysis is desired enter an empty list: `[ ]`. Otherwise, enter the name of the analysis functions separated by commas. Each name must be put in quotation marks, e.g. `['plot_model', 'landscape']`. A complete overview can be found in **Analysis method overview**. For the first run of a custom model, `['plot_model']` is recommended.
20. **Optional: enter customization options** – The behavior of the analysis functions can be customized. Possible options are included in the **Analysis method overview**. These values should be entered as pairs, similar to fit parameters (see **Intermezzo: Python types**). When opening the main script for the first time, the example values are formatted with each pair on a single line for readability, this is not required however.
21. **Run the framework** – When you are satisfied with the input options, run the script in order to start the framework. This can be done using the green arrow in Spyder or by pressing F5. The model will display the current activities in the output window (bottom right in the standard Spyder layout). Any generated images can be seen by pressing the 'plots' tab just above this window.
22. **Check point estimates** – Carefully inspect the determined point estimates. If the estimates differ significantly from expected values this could mean an error was made in the config or in the entered options. The analysis '*plot_model*' can help with a visual representation. **CRITICAL** It is also possible that the *system description* is incorrect. Perhaps an effect that was deemed negligible should be added after all, or the components of the system react in a different manor than was expected initially. A different *custom model* can be created to test different *system descriptions.* Go back to the appropriate step to generate the file and then return to this step.
23. **Perform additional analysis / change input** – The above steps can be repeated in order to test different models, different inputs (e.g. different compounds) or change the analysis functions to execute.

**Model builder rules**

When opening the model builder for the first time, notice how the example reactions are each defined on a single line. When entering or editing equations, make sure that the triple quotes surrounding the lines remain. An equation is defined as followed:

$$A + B = AB; Kd1$$

The left side contains one or more species that join to form the complex on the right side of the equality sign. Note that the right side is limited to a single species, use intermediate steps if this is not the case. The semicolon is followed by the equilibrium dissociation constant. A conformational change can be entered by having a single specie on both sides of the equality. The above example would result in the following equilibrium concentrations:

$$\frac{[A] * [B]}{Kd1} = [AB]$$

Each specie name should also follow a number of rules. The name can contain only alpha-numerical characters and **CRITICAL** must start with a capital letter. When multiple capital letters are present, the model builder will assume that it is a complex of multiple smaller species. e.g.: 'A', 'B', 'Enzyme', 'Q8' are all valid specie names. 'AB', 'EnzymeB', 'CaDaBa' are all valid names for the complexes of the species of A + B, Enzyme + B and Ca + Da + Ba respectively.

**CRITICAL** Note that due to these rules, the input AR + A will result in a different output than Ar + A. If multi-character specie names (e.g. Enzyme) are used, make sure that each component name is unique and not a subset of another name. E.g., defining both the specie 'Partner' and 'P' will result in errors to prevent unintended behavior.

These rules are in place in order to automatically fix common model definition mistakes. For example, in most cases when defining the following equations:

```
system_description = ("""
AB + C = ABC; Kd1
AC + B = ACB; Kd1
""")
```

the same complex $ABC$ should be formed by different routes, not two different complexes. Adhering to the above rules will allow the model builder to automatically correct these errors by always sorting the building blocks in the same manor. **CRITICAL** If this behavior is unwanted, instead use unique names containing only a single capital letter (e.g. $Abc$ and $Acb$).

Equilibrium constants can be defined as single constants or the product / ratio of multiple constants. Constant names must also consist of alpha-numerical characters and are case sensitive. They do not require capital letters and are not ordered automatically (as specie names are). Implicit multiplication, e.g. 2X, is not recognized and should be entered explicitly as 2 * X. Examples of rate definitions: 'Alpha', '2 * Kd1', 'Kd2 / alpha'.

### 2. Analysis method overview

| | |
|---|---|
| **plot_model** | **Plot the original data as points and the model prediction as a line based on the determined point estimates or the current parameter values in case no parameters have been fitted.** |
| ***Optional arguments:*** | |
| **model_plot_min** | float: The minimum titrate concentration to plot. *Default to all points* |
| **model_plot_max** | float: The maximum titrate concentration to plot. *Default to all points* |
| **normalise** | Boolean: If True will normalize all data in the plot between 0-1. Normalization factor is determined on a per condition basis. *Default False* |

| | |
|---|---|
| **Landscape** | **Plots a mean-squared error landscape for the fit parameters, displaying how the model error changes when the value of the parameters is moved from the determined point estimate. Line graph in case of only one fit parameter. In the case of two or more, will display a landscape using the first two parameters by default.** |
| ***Optional arguments:*** | |
| **landscape_parameters** | list of strings: The parameters to plot a landscape for. *Uses the fit parameters as default.* |
| **landscape_1_range** | 1d array-like: The points to plot for the first parameter, use of np.geomspace is recommended here. *The default range extends from 0.01 – 100 x the current parameter value.* |
| **landscape_2_range** | 1d array-like: The points to plot for the second parameter, if at least two landscape parameters are given. Use of np.geomspace is recommended here. *The default range extends from 0.01 – 100 x the current parameter value.* |

| | |
|---|---|
| **outlier** | **Uses a Doornbos procedure to attempt to detect outliers within technical repeats. Will only detect suspects, removal needs to be performed manually.** |
| ***Optional arguments:*** | |
| **write_output** | Boolean: Whether to create output files containing the suspect outliers. *The default is True.* |
| **skip_warnings** | Boolean: If True, will not print warnings about sets for which the Doornbos procedure is not defined. *The default is False.* |

| | |
|---|---|
| **plot_concentrations** | **Shows how the concentrations of all species in the system change over the range of titrate values. This function will use the current parameter values to determine concentrations.** |
| ***Optional arguments:*** | |
| **plot_species** | 'all' or 'components' or 'complexes' or list of strings: Which species to plot. The free titrate concentration is not plotted. *The default is 'all'*<br>'all' will show all species except for the free titrate.<br>'components': will plot all components, marked in the *custom model* file as independent species.<br>'complexes': will plot all complexes, marked in the *custom model* file as dependent species.<br>List of strings: Case sensitive, plot only species contained within the list. |

| | |
|---|---|
| **concentrations_start** | float: The lowest titrate concentration to plot. *The default is the lowest concentration in system.data.* |
| **concentrations_stop** | float: The highest titrate concentration to plot. *The default is the highest concentration in system.data.* |

| | |
|---|---|
| **confidence_interval** | **Determines a confidence interval for each fit parameter using the selected method, by default a non-parametric, bias corrected bootstrap approach.**<br>**The bootstrap method requires a large number of samples (recommend 1000 +) for accurate results.** |
| ***Optional arguments:*** | |
| **confidence_method** | 'bootstrap': Currently the only method supported. |
| **confidence_repeats** | integer: The number of repeats the confidence method should perform. |

| | |
|---|---|
| **parameter_sensitivity** | **Performs local parameter sensitivity analysis(1). The default m-function measures the change in total model error with a perturbation of +50% for each parameter in turn. See** Custom M functions **for details.** |
| ***Optional arguments:*** | |
| **sensitivity_perturbation** | float: The factor change in parameter value. Positive values are increase, negative values decrease, e.g. 0.5 will increase each parameter by 50%. *The default is 0.5.* |
| **sensitivity_state** | 'solution' or 'current': Which parameter values to use as reference. The currently assigned parameter values or the values determined during the fitting procedure (system.solution). *The default is 'current'.* |
| **sensitivity_parameters** | list of strings: Which parameters to test for sensitivity. *The default is the fit parameters.* |
| **m_function** | Python function: Used to define custom M-function. The function should take a single argument, the system object. *The default is the squared error function.* |

| | |
|---|---|
| **residuals** | **Displays the error between model prediction and actual measurement for each data point, centered around zero.** |
| ***Optional arguments:*** | None |

| | |
|---|---|
| **range_solver** | **Solve the system from a range of initial guess values, using a Latin hypercube approach to determine specific guess combinations. Displays all determined values sorted by the squared error.** |
| ***Optional arguments:*** | |
| **range_fit_parameters** | dictionary of string: tuple pairs. Each key represents a fit parameter, the tuple should contain the min and max expected values of that parameter / the boundaries of the Latin hypercube. *Required.* |
| **range_n** | integer: The number of strata to create within each range. *The default is 10.* |
| **log_scale** | Boolean: If True will use a log scale to determine strata. Alternatively use a regular linear scale hypercube sampling method. *The default is True.* |

### 3. Case I: Binding between 14-3-3 protein and partner protein, step-by-step guide

**System description (step 1)**

Here it is assumed that the user has read the *framework* procedure. This section primarily focuses on the specific values entered.

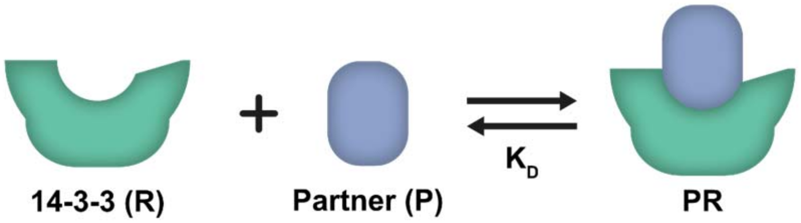

Figure S3: **System description for the introductory case**. The components R and P can complex to form PR. See the main text for further details.

The first system consists of two components that can complex together. A schematic overview of the *system description* is given in Figure S3. Within the description, a name is chosen for each specie and all the possible complexes are defined together with the dissociation constants. Writing the given system in the format expected by the model builder (see protocol / model builder rules) results in:

$$R + P = PR; Kd$$

**Model builder (steps 2-8)**

For this system, fluorescence anisotropy measurements were performed, where the partner ('P') is fluorescently labelled. In order to generate the correct data function, we use the 'anisotropy' function. Combined with the system description, the model builder is set up with the following values (note that the name does not influence the *custom model* and should be chosen for your convenience):

```
# model builder.py
# Case I
name = 'introduction'

system_description = ("""
R + P = PR; Kd
""")

data_function = "anisotropy"
labelled = "P"
custom_input = ""
```

After running the script, the *model builder* should report the following final equations:

$$(R - R\_tot + P * R/Kd) \,/\, R\_tot = 0$$

$$(P - P\_tot + P * R/Kd) \,/\, P\_tot = 0$$

After inspecting the output information, the 'introduction.py' file is moved from the './output' folder to the './model' folder.

**Experimental data and config file (steps 9-11)**

The experimental data gathered for this system is available in dataset S1. We chose to create a new input folder './input_introduction' for this system and place the excel file therein. The *config generator* creates the specification 'config.ini' file. The final input is:

```
# config generator.py
# Case I
input_file = ' input_introduction/dataset S1.xlsx'
```

This will generate a *config.ini* file in the './output' folder. Move the file to the './input_introduction' folder.

The next step is specifying the known concentrations of the *components*. The *model builder* reported the components in this system as `'R'` and `'P'`. While the total titrate concentration (`'R_tot'`) is contained in the input data, the total concentration of `'P'` will need to be defined in the *config.ini* file. A fixed partner concentration of 10 nM in all experiments can be specified by adding the line `P_tot = 10E-9` under the default section.

```
# config.ini
# Case I

[DEFAULT]
P_tot = 10E-9

[14-3-3 + Partner]
```

**Main script (steps 12-21)**
Within the main file, the name of the created model and the input folder are specified. In addition, the setup type 'separate' is chosen. This setup type creates an unique system for each experiment, but as the experimental data contains only a single experiment, only one system is created. Within the experiment, '`R`' was titrated. The model parameter that needs to be fitted is '`Kd`'. The initial guess for a parameter serves as the starting point for the numerical solver. Visual inspection of the experimental data suggests the $EC_{50}$ value to be around 5 µM and will serve as our initial guess. As the $K_D$ is the only parameter in this system there are no other known parameters to define. The only analysis that was performed for this system is '`plot_model`', this results in the following main file:

```
# main.py
# Case I
model = "introduction"
input_folder = "input_introduction/"

setup_type = "separate"
titrate = 'R_tot'

fit_parameters = {'Kd':5E-6}
known_parameters = {}

analysis = ['plot_model']
additional_arguments = { }
```

Running this file results in the following output, the model plot can be seen in the main text Figure 2C:

---14-3-3 + Partner---
Model parameters were determined using all available data points.
Best parameter estimates are:
Kd : 4.051e-06.
The root mean squared error with this fit is 1.56e+00

----------------------

**Check model (step 22)**

Root mean squared error is small compared to the experimental data values. The model prediction line correctly follows the data points. The determined parameter estimates correspond to previously determined values. It is therefore reasonable to conclude that this system is correctly modelled using the given *system description.*

**4. Case II: Protein-protein interaction (PPI) stabilization, step-by-step guide**
Here it is assumed that the user has read the *framework* procedure. This section primarily focuses on the specific values entered.

**System description (step 1)**

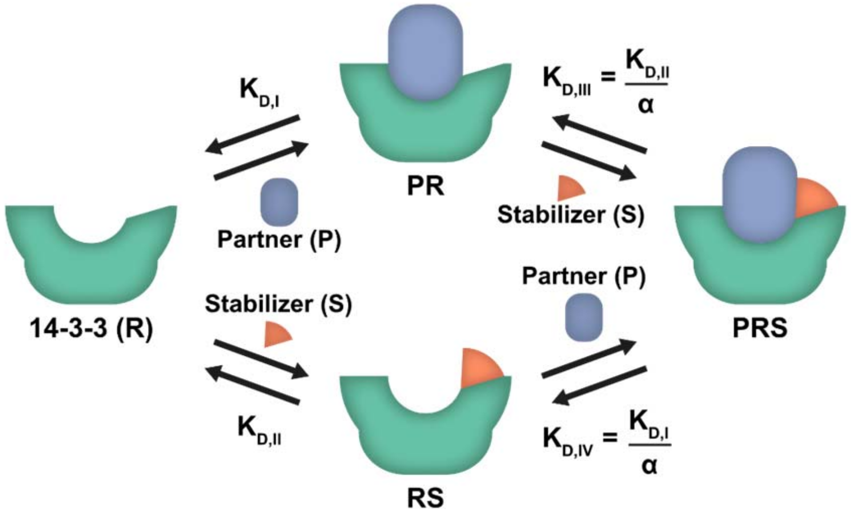

Figure S4: **System description for the protein-protein interaction stabilization case**. There are three components, P, R and S. This system contains two pathways to the final complex PRS. Either P or S can bind first to R. For an in-depth description see the main text.

The second system consists of three components that can complex together via two distinct pathways. The *system description* is given in Figure S4. Within the description, a name is chosen for each specie and all the possible complexes are defined together with the dissociation constants. Writing the given description in the format expected by the model builder (see protocol / model builder rules) leads to the following equations:

$$R + P = PR;\ Kd1$$

$$PR + S = PRS;\ Kd2\ /\ Alpha$$

$$R + S = RS;\ Kd2$$

$$RS + P = PRS;\ Kd1/\ Alpha$$

Note that this system has both multiple pathways resulting in the same complex and a thermodynamic cycle. Checks are present in the *model builder* which validate that the proposed system description is consistent, e.g. multiple pathways to the same complex consist of the same components and equilibrium constants and the product of all equilibrium constants in the cycle should result in 1.

**Model builder (steps 2-8)**
Similarly to the previous system, the experimental data consists of fluorescent anisotropy measurements (in 2 dimensions) with the partner labelled. Thus, the labelled component is `'P'`, and the data function is `'anisotropy'`. Combined with the system description, the model builder is set up with the following values:

```
# model builder.py
# Case II
name = 'PPI stabilization'
```

```
system_description = ("""
R + P = PR; Kd1
PR + S = PRS; Kd2 / Alpha
R + S = RS; Kd2
RS + P = PRS; Kd1/ Alpha
""")

data_function = "anisotropy"
labelled = "P"
custom_input = ""
```

After running the script, the *model builder* should report the following equations:

$$(Alpha * P * R * S/(Kd1 * Kd2) + S - S\_tot + R * S/Kd2) / S\_tot = 0$$

$$(Alpha * P * R * S/(Kd1 * Kd2) + R - R\_tot + R * S/Kd2 + P * R/Kd1) / R\_tot = 0$$

$$(Alpha * P * R * S/(Kd1 * Kd2) + P - P\_tot + P * R/Kd1) / P\_tot = 0$$

After inspecting the output information, the 'PPI stabilization.py' file is moved from the './output' folder to the './model' folder.

**Experimental data and config file (steps 9-11)**

The experimental data gathered for this system is available in **dataset S2**. The data contains 19 different experiments, one for each stabilizer concentration. We chose to create a new input folder './input_PPI' for this system and place the excel file therein. The config file is created using the *config generator.* Move the created 'config.ini' file from the './output' folder to the './input_PPI' folder.

As the titrate is 14-3-3 (R), the total concentrations of partner (*P_tot*) and stabilizer (*S_tot*) are defined in the config file. The total concentration of the partner is constant across all conditions and can be defined by adding `P_tot = 10E-9` to the default header. To define the stabilizer concentration in each experiment we add `S_tot = x` under each specific header, where `x` stands for the total concentration of stabilizer in that experiment.

```
# config.ini
# Case II

[DEFAULT]
P_tot = 10E-9

[ppi_1]
S_tot = 1E-9

[ppi_2]
S_tot = 2E-9

[ppi_3]
S_tot = 3E-9

[ppi_6]
S_tot = 6E-9

[ppi_12]
S_tot = 12E-9
```

```
[ppi_24]
S_tot = 24E-9

[ppi_49]
S_tot = 49E-9

[ppi_98]
S_tot = 98E-9

[ppi_195]
S_tot = 195E-9

[ppi_391]
S_tot = 391E-9

[ppi_781]
S_tot = 781E-9

[ppi_1563]
S_tot = 1563E-9

[ppi_3125]
S_tot = 3125E-9

[ppi_6250]
S_tot = 6250E-9

[ppi_12500]
S_tot = 12500E-9

[ppi_25000]
S_tot = 25000E-9

[ppi_50000]
S_tot = 50000E-9

[ppi_100000]
S_tot = 100000E-9

[ppi_200000]
S_tot = 200000E-9
```

**Main (steps 12-21)**

For this system, all the data should be fitted together to determine a single set of parameters, thus the 'Combined' setup type is chosen. One of the parameters has a known value as it was determined in the previous system ($K_{D,I}$: 4.05 µM), the other two parameters will be fitted by the *model solver*. Based on previous research, the values 66 µM(2) and 30(3) are used as initial guess for the $K_{D,II}$ and $\alpha$ parameter respectively. For this system the error landscape and the confidence interval were determined. In order to define the number of bootstrap samples, an additional argument is added resulting in the following main script:

```
# main.py
# Case II
model = 'PPI stabilization'
input_folder = "input_PPI/"
```

```
setup_type = 'combined'
titrate = 'R_tot'

fit_parameters = {'Kd2': 6.6E-5, 'Alpha': 30}
known_parameters = {'Kd1': 4.05E-6}

analysis = ['plot_model', 'landscape', 'confidence_interval']
additional_arguments = {
    'confidence_repeats': 2000}
```

Running this file results in the following output, the model plot is displayed in Figure S5.

---Combined system---
Model parameters were determined using all available data points.
Best parameter estimates are:
  Kd2 : 3.892e-04.
  Alpha : 1.335e+03.
The root mean squared error with this fit is 3.91e+00
---------------------

**Check model (step 22)**

Model prediction line deviates slightly at a few higher stabilizer concentrations. However, root mean squared error is small compared to the experimental data values and parameters agree with previously determined values. It is reasonable to assume that the *system description* is accurate, but it could prove interesting to investigate if additional effects are present at higher stabilizer concentrations.

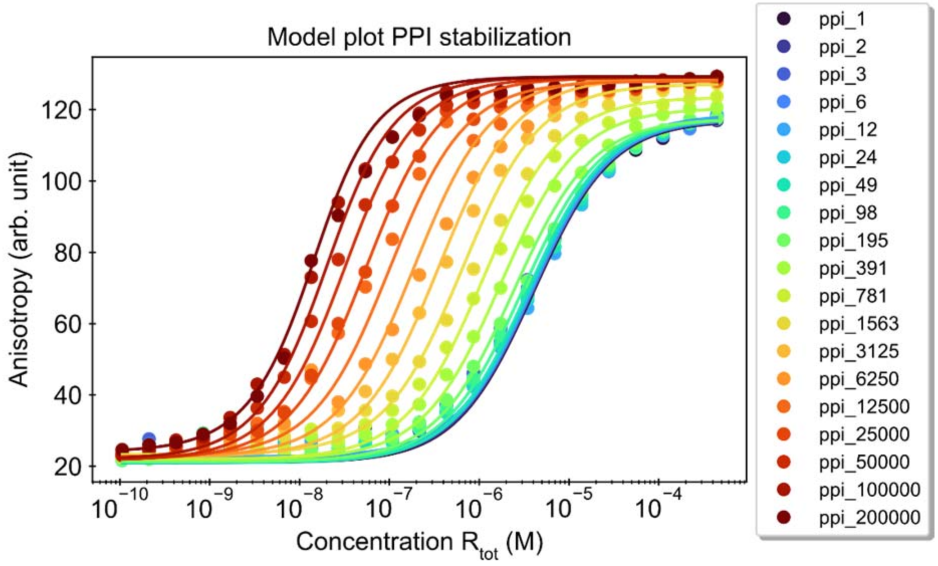

Figure S5: **'model_plot' for the protein-protein interaction stabilization case**. Fit parameters were given the following initial guess values: $K_{D,II}$: 66 µM, $\alpha$: 30. The value for $K_{D,I}$ was fixed at 4.05 µM. Determined estimates are $K_{D,II}$: 0.389 mM, α: 1.34 x $10^3$. The numbers in the labels indicate the concentration of the stabilizer in nM. The total concentration of partner was 10 nM in all measurements.

### 5. Bootstrap

The bootstrap method generates synthetic datasets based on random choice from the original data, in essence using the sample as a predictor for the real distribution(4–6). The parameter of interest is determined from these synthetic datasets in the same way as the original data. Repeating this procedure many times will result in a distribution for each parameter of interest from which a confidence interval can be determined. Within the *framework*, a bias corrected bootstrap method is used to determine confidence intervals for the fitted parameters.

The bootstrap method can be considered an application of Bayes' theorem(4) and thus the determined interval (intuitively) can be interpreted as the range of likely values. This is in contrast to a 95% confidence interval based on frequentist statistics. In those methods, the percentage refers to the confidence in the *method* used, rather than expressing the confidence in the *determined values*, even though they are commonly misinterpreted as such (7, 8). Another advantage of the bootstrap method is that the determined intervals are not required to be symmetrical by definition and any asymmetry in the samples is taken into account when determining the interval.

While the bootstrap method can take a significant amount of time, a large number of repeats is necessary for accurate estimates because of the random nature of the procedure. Bootstrapping with 2000 or more samples is common practice. The bootstrap results for the second case can be seen in Figure S6. Another fact to consider when using the bootstrap method is that it is sensitive to small sample sizes. Ten measurement points should be taken as the absolute minimum(9) with more measurements resulting in a better approximation. Having multiple independent measurements will improve the results by reducing any bias that might be introduced by the experimental setup while also increasing the number of data points. As the implementation for the *framework* is aimed at a wide range of possible experimental setups and usage scenarios, no specialized sampling methods are performed.

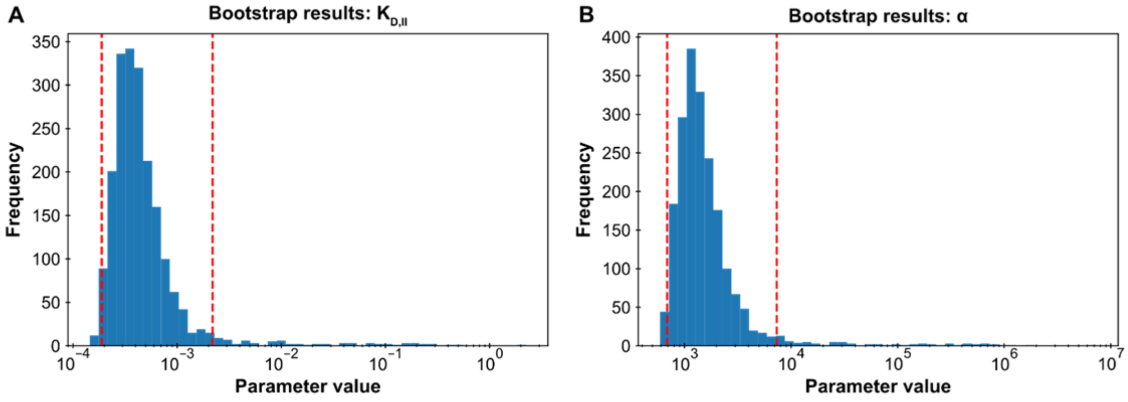

Figure S6: **Bootstrap results for the protein-protein interaction stabilization case.** The confidence interval analysis function was performed with 2000 repeats. The analysis creates a histogram of all determined values. The red lines indicate the determined 95% confidence interval. Both figures show a right-tailed log-normal distribution. **A:** $K_{D,II}$ parameter. **B**: $\alpha$ parameter.

### 6. Case III: Ratiometric Plug-and-Play Immunodiagnostics platform (RAPPID), step-by-step guide

Here it is assumed that the user has read the *framework* procedure. This section primarily focuses on the specific values entered.

#### System description (step 1)

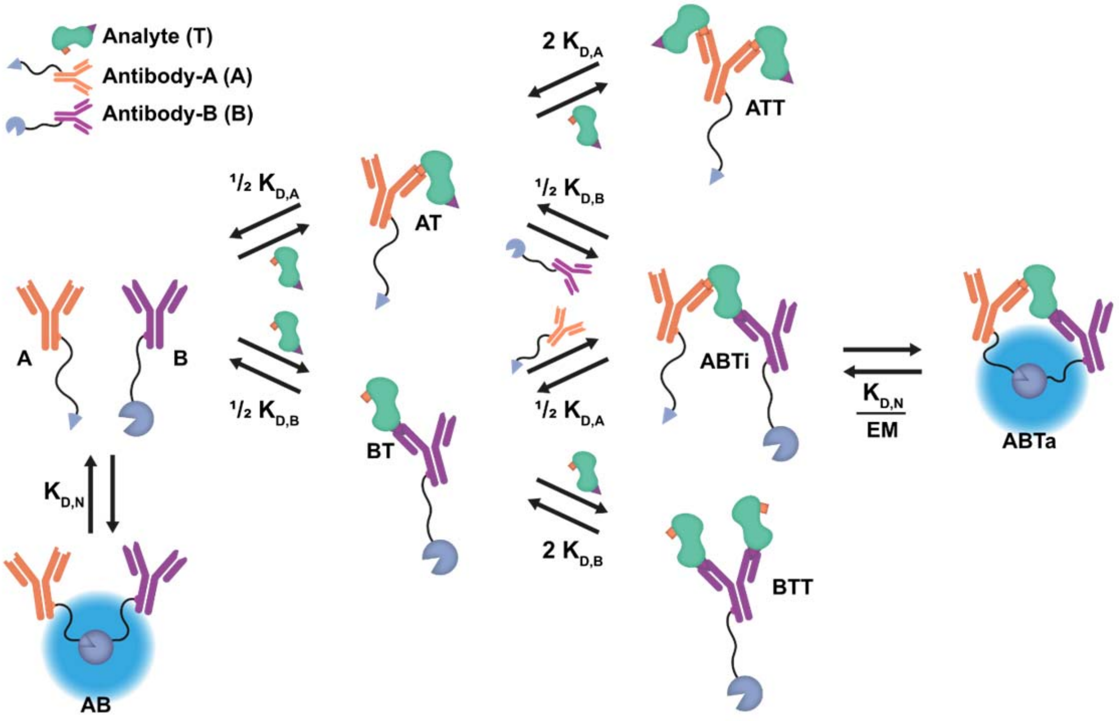

Figure S7: **System description for the RAPPID system**. This system contains three components, the Antibodies A and B and the analyte. It is described by a total of eight reversible reactions. For an in-depth description of the system, see the main text.

The final system consists of three components and each has multiple binding sites so multiple interactions are possible. The *system description* is given in Figure S7. Within the description, a name is chosen for each specie and all complexes are defined together with the dissociation constants. Writing the given description in the format expected by the model builder (see protocol / model builder rules) leads to the following equations:

$$A + B = AB;\ KdN$$

$$A + T = AT;\ 1/2 * KdA$$

$$B + T = BT;\ 1/2 * KdB$$

$$TB + T = BTT;\ 2 * KdB$$

$$TB + A = ABTi;\ 1/2 * KdA$$

$$TA + T = ATT;\ 2 * KdA$$

$$TA + B = ABTi;\ 1/2 * KdB$$

$$ABTi = ABTa;\ KdN\ /\ EM$$

#### Model builder (steps 2-8)

For this system, the experimental data consists of luminescence measurements. The relationship between the data values and the concentrations in the system is not defined in any of the standard data functions, so the 'custom' data function is chosen. The relationship is defined as the concentration of the active sandwich immunocomplex (ABTa) times some unknown scaling factor (see main text). The model builder is first executed without entering the custom input string. While the *custom model* cannot be finalized like this, the determined species can be inspected nevertheless. The name of the active complex is copied from the determined species and the custom function '`ABTa * Scaling`' is entered. See the **Custom data functions** section why this particular workflow is recommended. Combined with the system description, the model builder is set up with the following final values:

```
# model builder.py
# Case III
name = 'RAPPID'

system_description = ("""
A + B = AB; KdN
A + T = AT; 1/2*KdA
B + T = BT; 1/2*KdB
TB + T = BTT; 2*KdB
TB + A = ABTi; 1/2*KdA
TA + T = ATT; 2*KdA
TA + B = ABTi; 1/2*KdB
ABTi = ABTa; KdN / EM
""")

data_function = 'custom'
labelled = ''
custom_input = 'ABTa * Scaling'
```

After running the script, the *model builder* should report the following equations:

$$(4*A*B*EM*T/(KdA*KdB*KdN) + A*B/KdN + 4*A*B*T/(KdA*KdB) + A + 2*A*T/KdA + A*T**2/KdA**2 - A\_tot)\,/\,A\_tot = 0$$

$$(4*A*B*EM*T/(KdA*KdB*KdN) + 4*A*B*T/(KdA*KdB) + 2*A*T/KdA + 2*A*T**2/KdA**2 + 2*B*T/KdB + 2*B*T**2/KdB**2 + T - T\_tot)\,/\,T\_tot = 0$$

$$(4*A*B*EM*T/(KdA*KdB*KdN) + A*B/KdN + 4*A*B*T/(KdA*KdB) + B + 2*B*T/KdB + B*T**2/KdB**2 - B\_tot)\,/\,B\_tot = 0$$

The custom data function is also reported:

$$ABTa * Scaling$$

After inspecting the output information, the 'RAPPID.py' file is moved from the './output' folder to the './model' folder.

**Experimental data and config file (steps 9-11)**

The experimental data gathered for this system is available in **dataset S3**. The data contains a single experiment. A new input folder, './input_RAPPID' is created for this system and the excel file is placed therein. The config file is created using the *config generator.* After running the script, the 'config.ini' file is moved from the './output' folder to the './input_RAPPID' folder.

The titrate for this system is the analyte (T). The total concentrations of the A and B antibodies are therefore defined in the 'config.ini' file. The default header is used in order to reuse the config file for the analysis type system created later. While in luminescence measurements a substrate is converted in order to generate the signal, this specie is not directly modeled and can be assumed to be present in excess during the measurement and thus does not need to be defined in the config file.

```
# config.ini
# Case III

[DEFAULT]
A_tot = 1E-9
B_tot = 1E-9

[cTnI]
```

**Main (steps 12-21)**

For this system there are two main scripts. The first one is used to fit the cTnI data, the second is used to investigate the concentration plots. For the first script there are three fit parameters and two known parameters. Both the values for the known parameters as well as the initial guess values for the fit parameters are taken from the RAPPID publication.

```
# main.py
# Case III – RAPPID model comparison
model = 'RAPPID'
input_folder = "input_RAPPID/"

setup_type = 'separate'
titrate = 'T_tot'

fit_parameters = {'Scaling':1E17,'KdA':10E-9,'KdB':10E-9}
known_parameters = {'KdN': 2.5E-6, 'EM': 10E-6}

analysis = ['plot_model', 'parameter_sensitivity']
additional_arguments = {
    'sensitivity_parameters': ['KdN', 'EM', 'KdA', 'KdB']}
```

Running this file results in the following output, the model plot is available in Figure S8, the parameter sensitivity analysis in main text Figure 4B.

---cTnI---
Model parameters were determined using all available data points.
Best parameter estimates are:
  Scaling : 5.531e+17.
  KdA : 5.333e-07.
  KdB : 1.531e-08.
The root mean squared error with this fit is 1.28e+05
----------

**Check model (step 22)**

Compared to the experimental data values, the root mean squared error is small. Determined values are identical to those determined in RAPPID model. However, few experimental data points compared to the number of fit parameters.

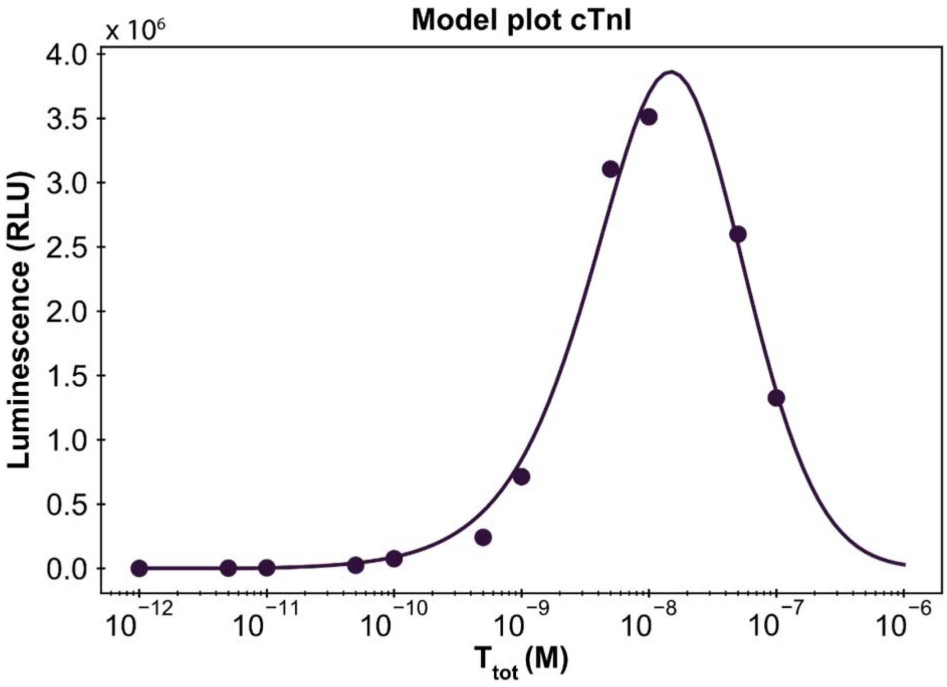

Figure S8: **Framework model plot for the RAPPID case.** Fit parameters were given initial guess values: *Scaling*: 1 x $^{17}$, $K_{D,A}$: 10 nM and $K_{D,B}$: 10 nM. $K_{D,N}$ and *EM* values were kept fixed at 2.5 µM and 10 µM respectively with $[A]_{tot}$: 1 nM and $[B]_{tot}$: 1 nM. The same initial conditions as were used in the RAPPID manuscript (see main text). Determined point estimates: *Scaling*: 5.53 x $10^{17}$, $K_{D,A}$: 533 nM, $K_{D,B}$: 15.3 nM, identical to the estimates determined by the model in the RAPPID publication. The option 'model_plot_max': 1E-6 was used to extend the plotting interval.

#### Main – concentrations plot

Note that this concentrations plot uses an *EM* value of 100 µM to clearly visualize the bell-shape in active complex concentration compared to the other specie concentrations. The results can be seen in main text Figure 4C.

```
# main.py
# Case III - No experimental data analysis
model = 'RAPPID'
input_folder = "input_RAPPID/"

setup_type = 'analysis'
titrate = 'T_tot'

fit_parameters = {}
known_parameters = {'KdN': 2.5E-6, 'EM': 100E-6, 'Scaling':1E17,
'KdA':10E-9,'KdB':15E-9}

analysis = ['plot_concentrations']
additional_arguments = {
    'plot_species': ['A', 'B', 'AB', 'ABTi', 'ABTa', 'AT', 'ATT'],
    'min_concentration': 5E-12,
    'max_concentration': 1E-6}
```

### 7. Custom data functions

During the fitting procedure, the data function predicts the experimental value corresponding to the current system specie concentrations. This prediction is compared to the experimental data and parameters are adjusted as needed. While several default experiment types are supported by the model builder, some systems and experiments will have a unique relation which requires a custom definition. See the RAPPID case in the main text for an example. **CRITICAL** It is important to note that because of the diversity of custom input equations, it is impossible to check for all possible errors, it is therefore paramount the user carefully reviews the processed equations and names of species (see below).

A custom definition can be created by choosing the 'custom' data function during the model builder. Below the data mode, a custom input can be given that defines the relationship between the system and the experimental data values as a string. The following rules apply.

First, any basic mathematical operation can be used to define the relationship, including powers, multiplication, division, brackets, subtraction and addition. Second, numbers can be entered separated by mathematical operators. No implicit multiplication, e.g. use 2 * x instead of 2x. Third, any unknown terms are transformed into new variables for the model. Fourth **CRITICAL** the names are case-sensitive and are not processed in the same way as the system description equations. In practice, this means they are not sorted based on building blocks (capital letters).

In order to get the concentration of a certain species in the system as part of the input equation, enter the name of that specie. Note that the name should be the same as processed version of that specie. It is therefore recommended to first run the model builder without a custom input (e.g. `""`), to determine if the model equations are correct. Once confirmed, the names of all the species will be visible at the top of the output provided by the model builder. These names can be used to get the concentrations of specific species. **CRITICAL** any other names will be interpreted as new variables!

An example: the system is described by the following input equation:

```
system_description = ("""
A + B = BA; Kd
""")
```

Note that the specie "BA" will be converted to "AB" during the model builder (see **Model builder rules** for details.) For the sake of example, we assume that the experimental data collected relates to the concentration of the complex "BA" times some constant. Entering the custom input:

`"BA * constant"`

will results in an error by the *model builder* as no specie concentrations were detected in the custom function. After first checking the model builder output, the following species are found in the input equations: A, B, AB. The correct custom input to describe the situation is therefore:

`"AB * constant"`

The final, correct input then becomes:

```
# model builder.py
system_description = ("""
A + B = BA; Kd """)
data_function = 'custom'
labelled = ''
custom_input = 'AB * constant'
```

**8. Custom M functions**

By default, the local parameter sensitivity function uses the system error function as the tracked value when performing the sensitivity analysis. By using the 'sensitivity_m_function' additional argument however, a M-function of choice can be inserted into the analysis. Writing custom functions does require familiarity with the Python programming language. The custom function should take a system object as parameter and should return a single value as output. The provided default analyses within the *framework* all return a pandas dataframe with the calculated values after completing the analysis. This fact can be leveraged to repurpose these analyses.

As an example, a M-function based on the maximum amount of active complex formation (ABTa) in the RAPPID *custom model* is created by repurposing the concentrations plot. The function takes a system object as parameter and calls the plot concentrations analysis on it, limiting the species only to the active complex and saving the returned value (the dataframe). The maximum value in this dataframe is returned as the M-function value.

```
# Custom M function definition
def max_concentration_m_function(system):
    df = src.post_process.plot_concentrations(system, plot_species=['ABTa'])
    return df['ABTa'].max()
```

Performing the parameter sensitivity analysis on the RAPPID system with the custom M-function results in the graph shown in Figure S9. The *EM* parameter has the greatest influence on the maximum amount of active complex formed. Meanwhile, the binding constants all show negative values, indicating an inverse relationship. Decreasing the dissociation constant will increase the amount of active complex formed.

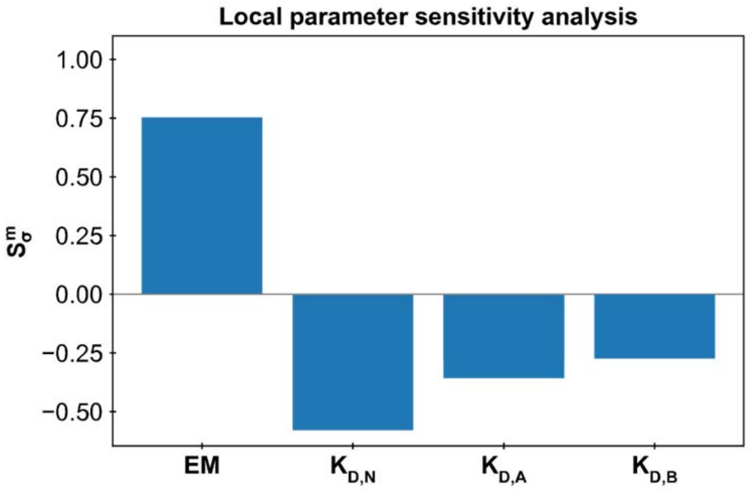

Figure S9**: Local parameter sensitivity analysis with custom M-function**. The M-function used measures the change in maximum active complex formed after an increase of 50% for each of the parameters. Known parameters were set to $K_{D,N}$: 2.5 µM, $K_{D,A}$: 15 nM, $K_{D,B}$: 10 nM, *EM*: 10 µM and *Scaling*: 1 x $10^{17}$ with no fit parameters. Both antibodies have a total concentration of 1 nM.

### 9. RAPPID model assumption analysis

The *system description* displayed in the main text, Figure 4A, is a direct representation of the model presented in the publication about the RAPPID system(10). Additional complexes can potentially form in this system which are not taken into account in the current description under the assumption that these form in negligible amounts. To assess if this assumption is valid, the current model can easily be extended and analyzed using the *framework.*

To do so, two additions will be made to the current *system description*. First of all, it is possible for either antibody (A / B) bound to the analyte (T) to form the active complex AB without forming the sandwich ternary complex. Secondly, it is possible that a second antibody binds to the ATT and BTT complexes. These complexes can then form an active complex consisting of four components. Alternatively, this state can be reached from the ABTa complex by binding an analyte on either side of the active complex. The extended system description can be seen in Figure S10.

For the added complexes, the order of the species matters (e.g. two T's bound to either A or B) and therefore unique names containing a single capital letter are used to identify each complex and differentiate the order of components within the complex. In addition, to prevent confusion with the A-antibody, the letters 'i' for inactive and 's' for signal/ active are used to represent the two states. The model builder is setup with the following equations. Note that the custom data function is also changed to include the four-component active complexes. The Tab and Tba complexes are considered part of the base signal together with the AB complex.

```
# model builder.py
# Case III – Extended RAPPID model
name = 'RAPPID - Extended'

system_description = ("""
A + B = AB; KdN
A + T = TA; 1/2*KdA
B + T = TB; 1/2*KdB
TB + T = TTB; 2*KdB
TB + A = ABTi; 1/2*KdA
TA + T = TTA; 2*KdA
TA + B = ABTi; 1/2*KdB
ABTi = ABTa; KdN / EM

TB + A = Tba; KdN
TA + B = Tab; KdN
TAT + B = Tatbi; 1/4*KdB
TBT + A = Tbtai; 1/4*KdA
Tatbi = Tatbs; KdN / EM
Tbtai = Tbtas; KdN / EM
ABTa + T = Tatbs; KdA
ABTa + T = Tbtas; KdB
""")

data_function = 'custom'
labelled = ''
custom_input = '(ABTa + Tatbs + Tbtas) * Scaling'
```

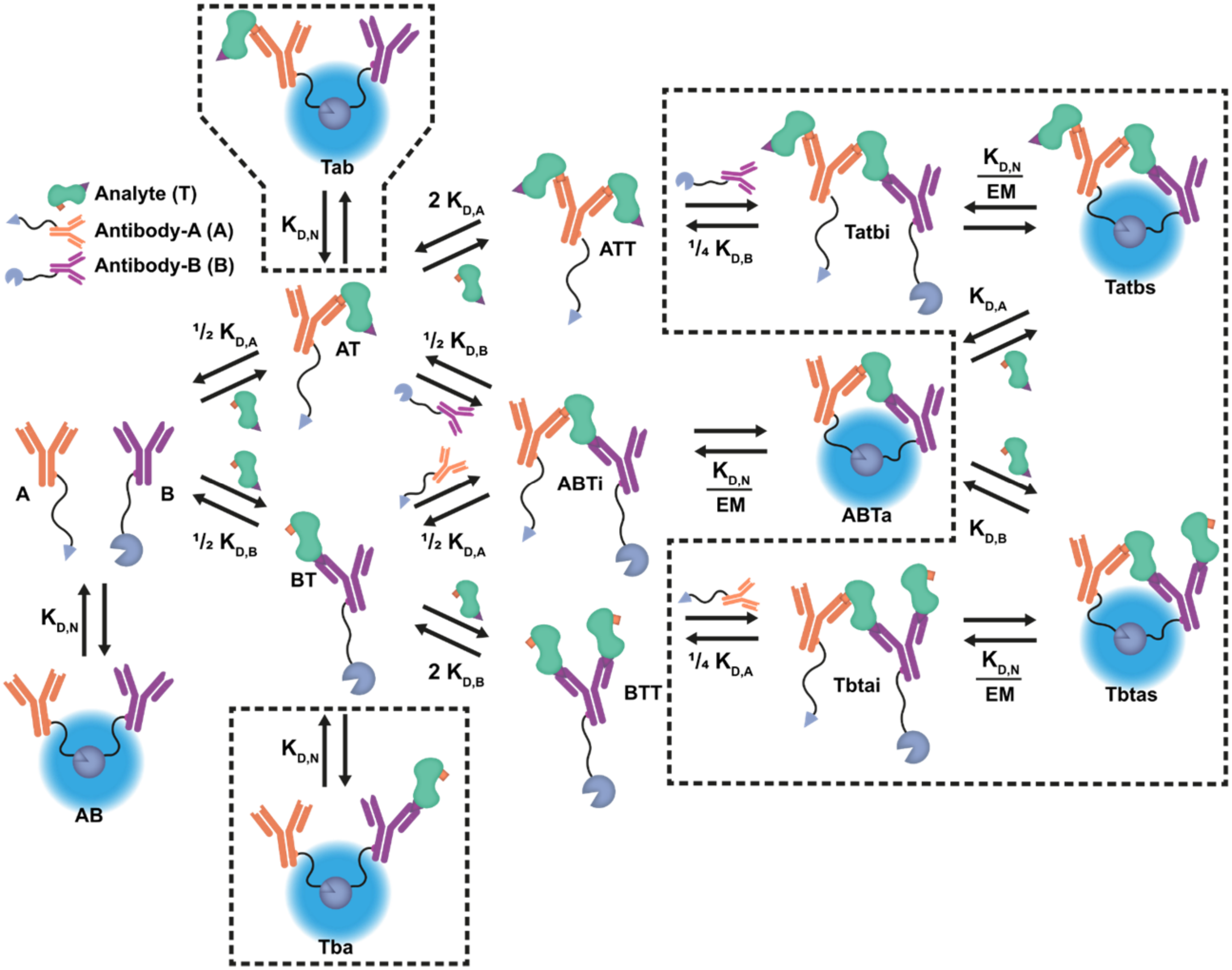

Figure S10: **RAPPID system extended system description.** Several complexes were added in order to test assumptions in the original system description. Both antibody-analyte complexes can now form the active NanoLuc complex with their respective partner which is still free in solution. In addition, quaternary complexes have been added to the system description. The additions to the original system description are marked with a dashed line.

First the formation of the Tba and Tab complexes is analyzed using the concentrations plot, using the previously determined point estimates as values for the parameters (Figure S11A). The maximum concentration reached for the Tba complex within the titration range is 0.183 pM. The maximum for Tab is about 1% of this concentration at 5.25 fM. These concentrations are indeed negligible, and it is therefore deemed safe to exclude them from the system description.

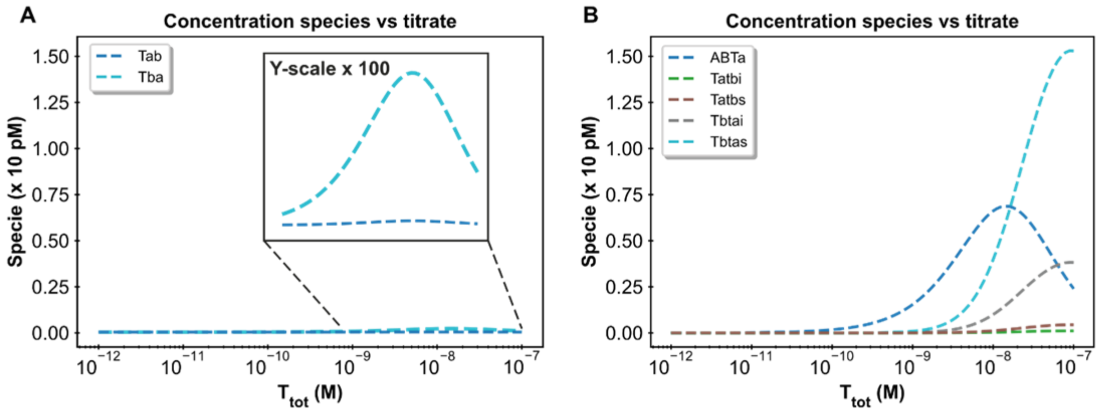

Figure S11 **Concentration plots for the extended RAPPID custom model:** Both images were generated based on the following fixed parameter values: $K_{D,N}$: 2.50 µM, $K_{D,A}$: 533 nM, $K_{D,B}$: 15.3 nM, *EM*: 10 µM, $[A]_{tot}$ = 1 nM and $[B]_{tot}$ = 1 nM **A:** Concentrations plot displaying Tab and Tba complex concentrations. **B:** Concentrations plot for the quaternary complex species compared to the active ternary ABTa complex.

Next, the formation of the quaternary active complexes was investigated, and the concentrations plot is again used with the previously determined point estimates (Figure S11B). In this case, it was found that the concentrations of the additional complexes within the titration range are not negligible and even surpass the concentration of the active complex at higher analyte concentrations. Fitting parameters to this extended *custom model* results in the following parameter estimates:

---cTnI---
Model parameters were determined using all available data points.
Best parameter estimates are:
    Scaling : 3.753e+16.
    KdA : 4.672e-08.
    KdB : 9.708e-09.
The root mean squared error with this fit is 1.29e+05
----------

These new parameter estimates confirm that the additional species cannot be ignored when modelling the system, as there is an order-of-magnitude difference between the estimates of both *custom models.* The size of this extended system makes manual derivation of the equilibrium equations extremely labor intensive, and this is where the *framework* excels, allowing easy extension of the existing system and the ability to check similar assumptions within minutes.

### 10. Custom analysis functions

The *framework* main script provides a 'façade' for the underlying code, providing an easy point of access for users less familiar with the Python programming language. The main script calls the src.driver.run function which validates the entered options and tries to locate the requested analyses. After all requested functions have been performed, the driver returns a list of all created systems to the main script. This procedure facilitates two options to add additional analyses to the *framework*.

The most direct method is using the list containing the returned system object(s) and writing code that performs additional operations on these objects. This could be in the form of a function or by just extending the main script after the call to the src.driver module. This method is ideal for one-off analyses or code that will only be used personally.

In order to fully integrate a new analysis into the *framework* however, a specific format should be followed. The driver module will attempt to find the analysis function requested by the user in the src.post_process module. Custom analysis should be added to this module in the form of a function with an all-lower-case name, using the '_' in order to improve readability if necessary (in accordance with the PEP style guide). The first and only positional argument excepted by the function must be a system object. All other arguments must be named arguments. The function should also contain a '**kwargs' argument as each analysis is offered all additional arguments entered in the main script. This structure is aimed at providing maximum ease-of-use for the users of the main script but does provide some additional hurdles for analysis writers. The analyses in the *framework* can be used as a style guide or example. As soon as the function is added to the post_process module, the analysis can be called by adding the function name to the desired analyses list.

It is recommended that each named argument starts with a part of the name of the analysis function to prevent frequent term overlap. As an example, the confidence interval function uses the 'confidence_repeats' argument instead of just 'repeats'. It is also recommended to use default values wherever possible to simplify the usage of the new analysis. Complex analysis can be contained in their own module and the function in the post_process module can then be used to redirect to this module. Finally, a decorator is available to add the 'write to file' functionality to new analysis functions. This functionality can be added to any function that returns a pandas dataframe (or other object that has a 'to_csv' method) by adding '@write_to_file' above the function definition. Once again, the analyses present in the post_process module can be used as an example.

### 11. Stress testing

Two types of extreme *system descriptions* were used to test the performance and capabilities of the *framework*. The first type uses linearly extending complexes. In these systems, component A binds to component B in order to form complex AB. Complex AB then binds to component C to form complex ABC, etc. In this approach, each individual equilibrium equation is simple but for a maximum complex size n, there are also n equilibrium equations that need to be solved simultaneously. In the second type, a combinatorial method is used. The system with the component A, B and C can form the complexes AB, AC, BC and ABC. Each additional component thus exponentially increases the number of complexes that can be formed while only increasing the number of equilibrium equations by one. In this setup both the number of components (n) and the maximum size of the complexes formed (m) can be changed in order to reach a specific number of complexes in the system.

For each method, a concentrations plot was created for the first component in the *custom model.* This will make sure a solution was found to the system of *equilibrium equations* for one hundred different titrate concentrations while keeping the comparison as fair as possible. The component concentrations were set to 15 M with random noise from a normal distribution ($\mu = 0, \sigma = 2{,}5$). Molar range concentrations were used to prevent the final complex concentrations (e.g. with 100 complexes) from becoming too small for the calculations. The time required to complete the calculations was tracked using the src.analysis.helpers timer. The results are displayed in Figure S12.

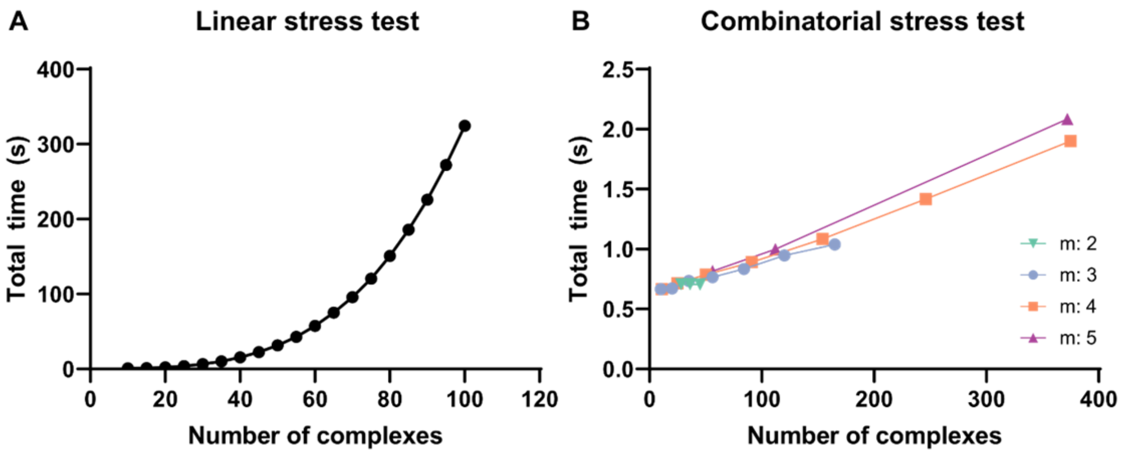

Figure S12: **Stress test results for the framework. A:** Computational time required for the linearly extending custom models. A quadratic increase in the required time can be observed with increasing component numbers. While the absolute time requirement is pc dependent, the relationship is not. **B:** Computational time required for the combinatorial custom models. Barely any increase in computational time is observed even though the total number of complexes is increased a 100-fold. The total number of complexes possible is dependent on both the number of components (n) in the system and the maximum number of components allowed in a single complex (m).

The *framework* was able to correctly determine equilibria conditions for all tested *custom models.* Comparing the two approaches, it becomes apparent that the total number of equilibrium equations, which is determined by the number of components in the *system description*, is the decisive factor in the total duration of the computations. Equilibrium conditions for one hundred different titrate concentrations for systems consisting of over 350 complexes were still determined within mere seconds. A linear relationship can also be observed between the total number of complexes and the required computational time. Meanwhile, increasing the number of components leads to a quadratic increase in computational time. While there is a clear increase in the time required, even systems of a hundred unique components could be solved by the *framework,* which is well beyond most typical experimental multi-component systems.
